# TNF-*α* disrupts the malate-aspartate shuttle, driving metabolic rewiring in iPSC-derived enteric neural lineages from Parkinson’s Disease patients

**DOI:** 10.1101/2025.03.25.644826

**Authors:** Bruno Ghirotto, Luís Eduardo Gonçalves, Vivien Ruder, Christina James, Elizaveta Gerasimova, Tania Rizo, Holger Wend, Michaela Farrell, Juan Atilio Gerez, Natalia Cecilia Prymaczok, Merel Kuijs, Maiia Shulman, Anne Hartebrodt, Iryna Prots, Arne Gessner, Friederike Zunke, Jürgen Winkler, David B. Blumenthal, Fabian J. Theis, Roland Riek, Claudia Günther, Markus Neurath, Pooja Gupta, Beate Winner

## Abstract

Gastrointestinal (GI) dysfunction emerges years before motor symptoms in Parkinson’s disease (PD), implicating the enteric nervous system (ENS) in early disease progression. However, the mechanisms linking the PD hallmark protein, α-synuclein (α-syn), to ENS dysfunction - and whether these mechanisms are influenced by inflammation - remains elusive. Using iPSC-derived enteric neural lineages from patients with α-syn triplications, we reveal that TNF-α increases mitochondrial-α-syn interactions, disrupts the malate-aspartate shuttle, and forces a metabolic shift toward glutamine oxidation. These alterations drive mitochondrial dysfunction, characterizing metabolic impairment under cytokine stress. Interestingly, targeting glutamate metabolism with Chicago Sky Blue 6B restores mitochondrial function, reversing TNF-α-driven metabolic disruption. Our findings position the ENS as a central player in PD pathogenesis, establishing a direct link between cytokines, α-syn accumulation, metabolic stress and mitochondrial dysfunction. By uncovering a previously unrecognized metabolic vulnerability in the ENS, we highlight its potential as a therapeutic target for early PD intervention.

## Introduction

Parkinson’s disease (PD) is the most frequent movement disorder in humans. The motor deficits - hypokinesia, rigidity, and tremor - are caused by dopaminergic neuronal loss in the substantia nigra^1^. However, non-motor symptoms, particularly those affecting the gastrointestinal (GI) tract, are increasingly recognized as early and clinically significant manifestations of PD^2,3^. Chronic constipation and abdominal bloating often precede motor deficits by years^4^, suggesting a role for the enteric nervous system (ENS) in early disease pathogenesis. The ENS, a vast network of over 500 million enteric neurons and glia within the bowel wall, regulates motility, secretion, and blood flow while also modulating intestinal immunity and interacting with the microbiota^5^. Its striking similarities to the central nervous system (CNS) regarding neuronal diversity and signaling suggest that ENS dysfunction could contribute to early GI symptoms in PD and influence disease progression.

Accordingly, Braak’s hypothesis suggests that PD pathology originates in the GI tract and spreads to the brain via the vagus nerve^6^. Central to this hypothesis is α-synuclein (α-syn), whose aggregation defines PD pathology^1^. While α-syn toxicity has been extensively studied in CNS^7^, its impact on the ENS remains poorly understood. Histopathological studies confirmed α-syn accumulation in enteric neurons of PD patients^8^, and animal models replicated this phenomenon, linking ENS dysfunction to GI symptoms^9,10^. Notably, injection of α-syn epitopes in humanized mice triggered intestinal inflammation, constipation, and enteric neuronal loss, demonstrating that α-syn can directly drive ENS dysfunction and GI pathology^11^.

Neuroimmune interactions, the crosstalk between neuronal and immune signaling, are increasingly recognized in PD, with cytokines playing a key role in α-syn-driven pathology^12-14^. We previously demonstrated that α-syn increases the susceptibility of iPSC-derived cortical neurons to cytokine-induced toxicity, particularly IL-17A^14^, suggesting that both α-syn accumulation and inflammatory stimuli contribute to disease pathology. In the CNS, proinflammatory cytokines exacerbate α-syn-induced mitochondrial dysfunction, oxidative stress, and synaptic impairments^1,13,14^, accelerating neurodegeneration. Similarly, α-syn has been implicated in intestinal inflammation^9,11,15,16^, yet most studies rely on animal models, which do not fully replicate human pathology. Given that both immune activation and mitochondrial dysfunction play essential roles in PD, metabolic dysregulation may act as a key intermediary linking these processes. In this sense, impaired energy metabolism and oxidative stress have been described as major contributors of PD pathogenesis^1,17^, with enteric neurons being particularly dependent on mitochondrial respiration^18^. Disruptions in metabolic homeostasis may amplify their vulnerability to inflammation, further aggravating ENS dysfunction. PD studies have demonstrated disease-induced alterations in mitochondrial respiration, amino acid metabolism, and bioenergetic substrate utilization^17,19^, but how these mechanisms contribute to enteric neurodegeneration remains unclear.

Studying α-syn pathology in the human ENS faces significant challenges. Unlike classical gastrointestinal diseases like inflammatory bowel diseases (IBD), GI specimens from PD patients are rarely available. ENS complexity and its unique microenvironment limit insights from histopathological studies, which confirmed α-syn accumulation^8^, but not its functional impact. Additionally, interspecies differences in animal models hinder the translation of findings to human disease^20^. Recent iPSC advances offer a promising platform for modeling α-syn pathology in both the CNS and ENS. iPSC-derived neurons enable patient-specific studies^21^, and protocols for generating enteric neural lineages (ENLs) from iPSCs^22^ have advanced Hirschsprung disease^23^. Nevertheless, this approach remains unexplored for PD.

To address these gaps, we utilized iPSC-based modeling of the ENS. Employing multiomics and functional analyses, we examined whether an increase in α-syn contributes to early PD pathology in the ENS and if this process is intensified by proinflammatory cytokines. We used iPSC-ENLs from PD patients with α-syn gene triplications (SNCA 3x) and demonstrated that SNCA 3x ENLs exhibit enhanced TNF-α susceptibility, leading to increased mitochondrial α-syn accumulation. This triggered TNF-induced metabolic reprogramming, impairing the malate-aspartate shuttle (MAS) and tricarboxylic acid (TCA) cycle, thereby increasing reliance on glutamine oxidation. These metabolic shifts impaired cellular energy metabolism, leading to increased enteric neuronal vulnerability. This could be rescued by targeting glutamate metabolism with Chicago-Sky-Blue 6B (CSB6), a competitive inhibitor of vesicular glutamate transporters. Our data highlight the iPSC-ENL platform as an invaluable tool for studying human synucleinopathies and underscore cytokines and metabolic dysfunction, specifically MAS impairment, as critical drivers of enteric pathology in early PD.

## Results

### iPSC-ENLs as a model for studying synucleinopathies

To study the enteric nervous system in synucleinopathies, we generated ENLs from three SNCA 3x iPSC lines and their isogenic controls (Iso)^24^.

The differentiation protocol starts from iPSCs and involves the induction of neural crest cells followed by enteric lineage specification to recapitulate key developmental pathways in vitro^22^. Within the differentiation process towards ENLs we assessed the expression of marker genes at three time points: iPSC at an early stage of ENC induction (day 6), a middle time point during ENL differentiation (day 40) and full differentiation at day 70 (Figure 1A). As expected, already at day 40 of ENL differentiation, Reverse Transcription Polymerase Chain Reaction (RT-qPCR) confirmed the presence of the neuronal marker *TUBB3* and enteric neuronal markers *ELAVL4, PHOX2B*, and *HOXB3* in both Iso and SNCA 3x lines (Figures 1B, S1B). Similarly, ENLs showed expression of the glial marker *GFAP* at day 70, with no group-level differences (Figures 1B, S1B). Immunostaining confirmed the presence of GFAP and the enteric neuron marker HuC/D in both groups, along with a specific increase in α-syn levels in SNCA 3x ENLs at day 70 (Figure 1C).

**Figure 1.**
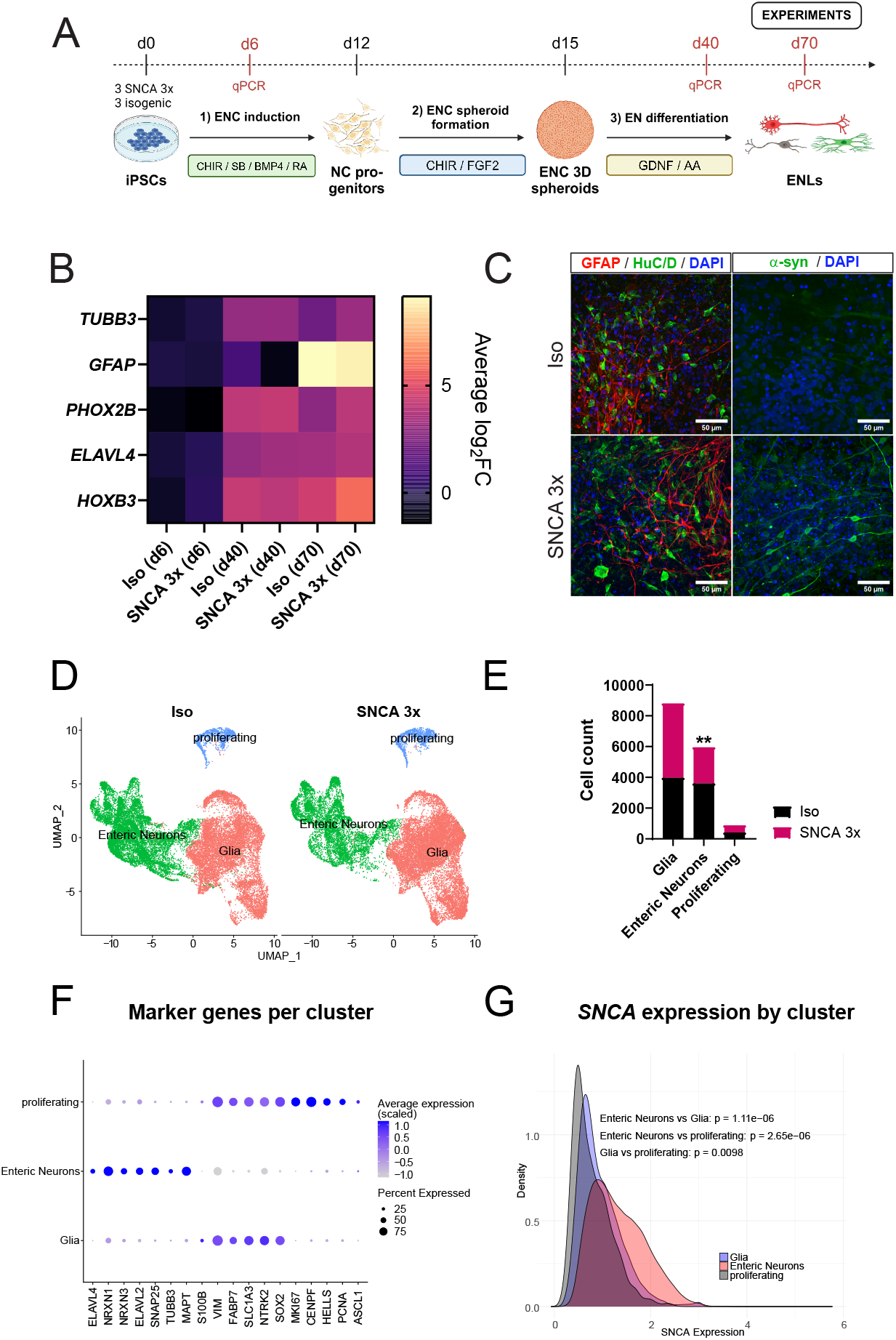
iPSC-ENLs as a model for studying synucleinopathies. **(A)** Paradigm illustrating the differentiation of ENLs from iPSCs. Generated using BioRender. **(B)** Heatmap showing gene expression of neuronal marker *TUBB3*, glial marker *GFAP*, and enteric neuronal markers *PHOX2B, ELAVL4*, and *HOXB3* in iPSC-derived ENLs at days 6, 40, and 70 of differentiation using RT-qPCR. Log2 fold change was calculated relative to the Iso group at day 6 of differentiation and averaged for n=3 biological replicates per group. See also Figure S1B. **(C)** Immunocytochemistry characterization of iPSC-derived ENLs at day 70 after start of differentiation. Left panels show the glial marker GFAP (red), the enteric neuronal marker HuC/D (green), and DAPI (blue) for cell nuclei staining. Right panels show α-syn (green, 2A7 antibody, Novus, Cat: NBP1-05194) and DAPI (blue) for cell nuclei staining. Upper panels represent the Iso group, and bottom panels the SNCA 3x group. **(D)** UMAP plots obtained from scRNA-seq analysis of Iso and SNCA 3x ENLs at day 70 after start of differentiation, separated by the clusters identified after annotation. **(E)** Fraction of cells per cluster identified in (D). n=3 biological replicates per group, **p<0.01 by unpaired two-tailed Student’s t-test. **(F)** Dotplot showing the average and percentage of expression of canonical marker genes for each cluster identified in (D). **(G)** Ridgeplot comparing the expression of *SNCA* between each cluster identified in (D). n=3 biological replicates per group, p-values calculated by unpaired two-tailed Student’s t-test.

Cellular diversity and gene expression patterns in iPSC-ENLs at day 70 of differentiation were explored using single-cell RNA sequencing (scRNAseq). Through clustering and subsequent annotation, we determined three primary cell populations: enteric neurons, glia, and proliferating cells (Figures 1D, S1C-D). Interestingly, SNCA 3x ENLs exhibited a lower proportion of enteric neurons compared to the Iso group (Figure 1E). Each cluster expressed its classically described canonical markers^22,25^ (Figure 1F). We also measured *SNCA* expression levels to assess α-syn in the ENS. Enteric neurons had the highest *SNCA* expression, followed by glia, and proliferating cells showed the lowest expression (Figure 1G). Between the two conditions, SNCA 3x glia exhibited higher *SNCA* expression compared to Iso (Figure S1E).

To highlight the broader relevance of our iPSC ENL model, we subclustered the scRNAseq enteric neurons and glia cell populations (Figures 1D and S1D) and compared them to the previously published single-cell atlas of human gut tissue^26^. We identified 14 neuronal subclusters and 13 glial subclusters (Figures 2A, 2E), with top markers associated with these subclusters depicted in Figure S2A-B and Table S2. Annotation using SingleR successfully mapped several distinct neuronal and glial subtypes to the reference human gut tissue (Figure S2C-F and Table S3). This included putative excitatory motor neuron 3 (PEMN3), expressing the D2 dopamine receptor and putative inhibitory motor neuron 3 (PIMN3), marked by adrenomedullin. The analysis also identified putative sensory neurons (PSN), which sense and respond to chemical and mechanical stimuli and interneurons (PIN), involved in intercellular communication (Figures S2C, S2E). Most enteric glia subclusters matched the Glia 1 subtype, linked to GDNF receptor *GFRA2* expression^26^ (Figures S2D, S2F).

**Figure 2.**
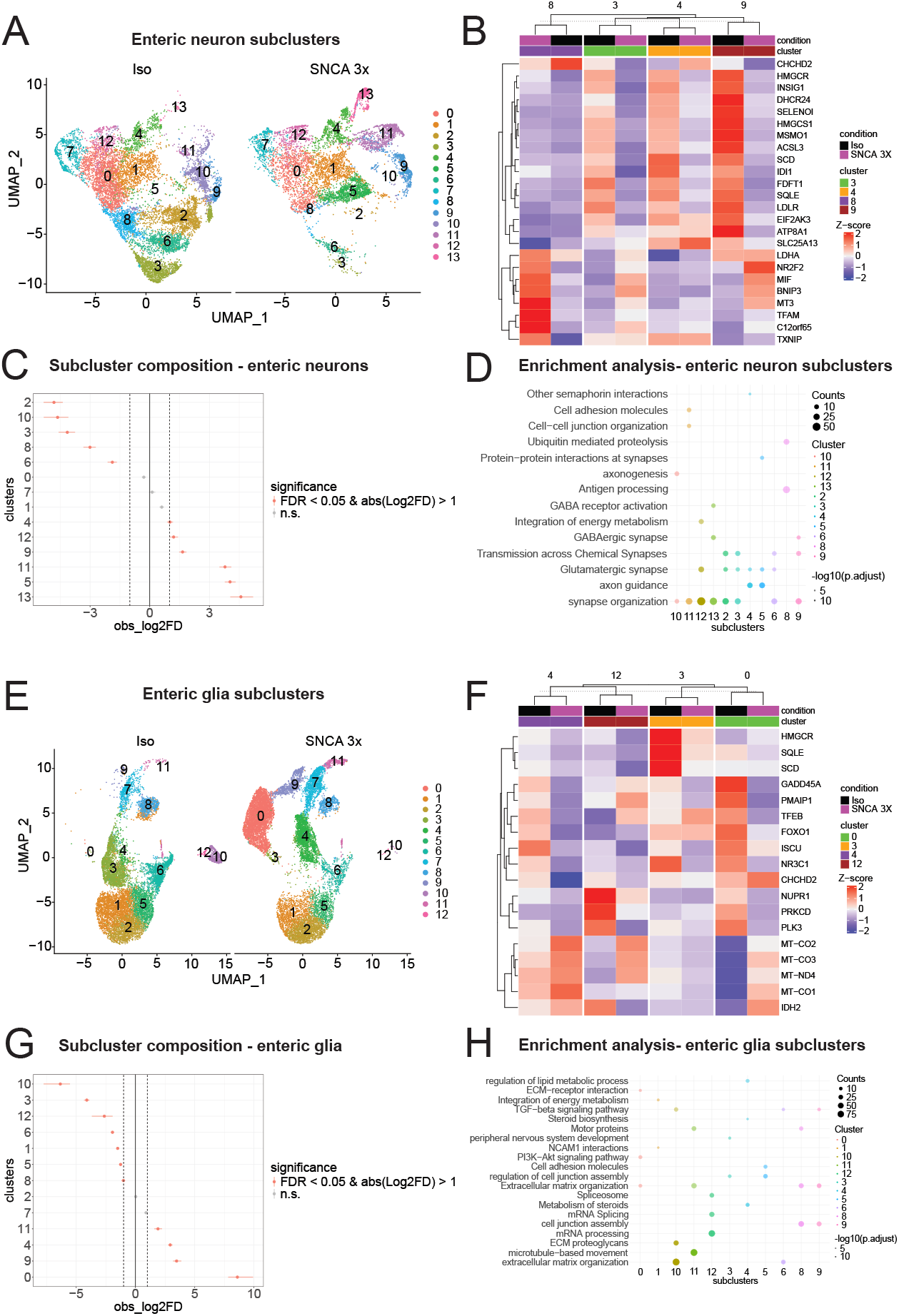
SNCA 3x drives mitochondrial dysfunction at transcriptional level and alters subpopulation abundance of enteric neurons and glia. **(A)** UMAP plots obtained from subclustering analysis of the enteric neurons (see figure 1D) of Iso and SNCA 3x ENLs at day 70 after start of differentiation. **(B)** Heatmap obtained from differential gene expression analysis of Iso vs SNCA 3x enteric neurons characterizing the expression of transcripts associated with the regulation of mitochondrial function. **(C)** Compositional analysis plot showing the significantly changed enteric neuron subclusters between Iso and SNCA 3x. FDR=False Discovery Rate (Benjamini-Hochberg correction); log2FD=log2Fold Difference. **(D)** Integrated enrichment analysis including Reactome, KEGG and Gene Ontology Biological Processes, Molecular Function and Cellular Component based on the enteric neuron subcluster markers depicted in Figure S2A and Table S2. See table S5. **(E)** UMAP plots obtained from subclustering analysis of the glia (see figure 1D) of Iso and SNCA 3x ENLs at day 70 after start of differentiation. **(F)** Heatmap obtained from differential gene expression analysis of Iso vs SNCA 3x enteric glia characterizing the expression of transcripts associated with the regulation of mitochondrial function. **(G)** Compositional analysis plot showing the significantly changed enteric glia subclusters between Iso and SNCA 3x. FDR=False Discovery Rate (Benjamini-Hochberg correction); log2FD=log2Fold Difference. **(H)** Integrated enrichment analysis including Reactome, KEGG and Gene Ontology Biological Processes, Molecular Function and Cellular Component based on the enteric glia subcluster markers depicted in figure S2B and table S2. See table S5.

Overall, these results indicate that both Iso and SNCA 3x iPSC lines differentiate effectively into ENLs, comprising a mixed culture of enteric neurons and glial cells, which recapitulate the cellular diversity of the human ENS. The observed α-syn expression underscores the model’s value for studying synucleinopathies and broader ENS research.

### SNCA 3x drives mitochondrial alterations at the transcriptional level and alters subpopulation abundance of enteric neurons and glia

We investigated whether increased α-syn alters gene expression and cellular abundance in enteric neuron and glial subpopulations using the scRNAseq data. Differential analyses revealed mitochondrial dysfunction, a PD hallmark^27^, as a key feature of SNCA 3x ENLs. The number of differentially expressed genes (DEGs) per subcluster is provided in supplementary table S4.

For the SNCA 3x enteric neurons, the subpopulations corresponding to subclusters 8, 3, 4, and 9 exhibited the highest changes in DEGs (Figure 2B, table S4). Specifically, subclusters 3, 4 and 9 showed downregulation of *HMGCR, DHCR24*, and *SQLE* (lipid homeostasis^28-30^), and *EIF2AK3* (stress response^31^) (Figure 2B, Table S4). In contrast, subcluster 8 upregulated *BNIP3* and *TXNIP* (mitochondrial stress^32,33^), *TFAM* (mitochondrial biogenesis34), and *LDHA* (metabolic reprogramming^35^) (Figure 2B, Table S4). Functional enrichment analysis of these subclusters revealed pathways related to synapse organization and glutamatergic signaling (subclusters 3, 4, 9) and antigen processing (subcluster 8), indicating that populations with metabolic alterations may exhibit synaptic dysfunction and immune-related stress responses (Figure 2D, Table S5). For SNCA 3x enteric glia, subclusters 4, 12, 3 and 0 exhibited notable changes in transcription profile (Figure 2F, Table S4). Subcluster 3, in particular, showed downregulation of *HMGCR, SQLE*, and *SCD*, indicative of disrupted cholesterol metabolism^28,30,36^ (Figure 2F, Table S4), which could impact membrane integrity and mitochondrial dynamics. Additionally, defects in lipid metabolism can disrupt β-oxidation, an essential mitochondrial process for ATP production, potentially worsening the metabolic vulnerability observed in SNCA 3x ENLs. Across all glial subclusters, *NUPR1, PRKCD*, and *PLK3* were downregulated, suggesting impaired oxidative stress response^37-39^ (Figure 2F, Table S4). Conversely, subclusters 0, 12, and 4 exhibited upregulation of oxidative phosphorylation genes *MT-CO2, MT-CO3, MT-ND4*, pointing to a potential compensatory increase in mitochondrial respiration^40^ (Figure 2F, Table S4).

Compositional analysis correlated with differential expression analysis findings (Figures 2C, 2G). The reduced abundance of neuronal subcluster 8 (Figure 2C) in SNCA 3x, combined with the upregulation of stress-adaptive and mitochondrial biogenesis genes (*BNIP3, TFAM, TXNIP*) (Figure 2B, Table S4), suggests selective vulnerability of this subpopulation towards stress. In SNCA 3x glia, subcluster 0 showed increased abundance (Figure 2G), in conjunction with upregulated oxidative phosphorylation genes (*MT-CO2, MT-CO3, MT-ND4*) (Figure 2F, table S4). On the contrary, depleted subclusters 3 and 12 (Figure 2G) were enriched for genes associated with the regulation of cell junctions and mRNA processing (Figure 2H, Table S5), processes essential for maintaining cellular homeostasis under stress.

These data suggest that SNCA 3x disrupts mitochondrial homeostasis in enteric neurons and glia at the transcriptional level, altering subpopulation dynamics and inducing metabolic stress-driven vulnerability.

### SNCA 3x drives basal mitochondrial dysfunction and alters enteric neuron-glia communication in iPSC-ENLs

Given the extensive transcriptional changes in mitochondrial-linked genes and the enrichment of metabolic pathways in SNCA 3x enteric neurons and glia (Figure 2) and their relevance to PD^1,17^, we wanted to validate the findings by analyzing different aspects of mitochondrial morphology. Using TOM20 immunostaining and automated image analysis, we assessed several mitochondrial parameters (Figures 3A, 3B). Interestingly, our data detected a significant decrease in mitochondrial count and area per cell in SNCA 3x ENLs (Figure 3B), suggestive of mitochondrial stress and mitophagy. Additionally, the number of branches and the branch junctions per cell were also significantly decreased in SNCA 3x ENLs (Figure 3B), indicating shortening of the mitochondrial network, which is a classical feature of mitochondrial fission. Next, we analyzed mitochondrial-linked gene expression along pseudotime (Figures 3C, 3D), using Slingshot^41^. Our results revealed that in SNCA 3x enteric neurons, genes controlling oxidative phosphorylation^40^, ATP production^40^ and mitochondrial biogenesis^42^ were downregulated (*ATP5F1B, COX4I1, NDUFA9, PPARGC1A*), while those controlling mitophagy^43^ (*PINK1*) and fission^44^ (*DNM1L*) were upregulated (Figure 3C). Conversely, in glial cells, *ATP5F1B, COX4I1*, and *NDUFA9* were upregulated along with *PINK1* and *DNM1L*, whereas *PPARGC1A* showed a transient peak before downregulation (Figure 3D). Complementarily, we performed a metabolic flux analysis focusing on mitochondrial function, which suggested an upregulation of Oxalosuccinate:NADP+ oxidoreductase in both SNCA 3x enteric neurons (Figure 3E) and glia (Figure 3F), indicating a shift toward NADPH production and highlighting potential compensatory metabolic adaptations to mitochondrial dysfunction.

**Figure 3.**
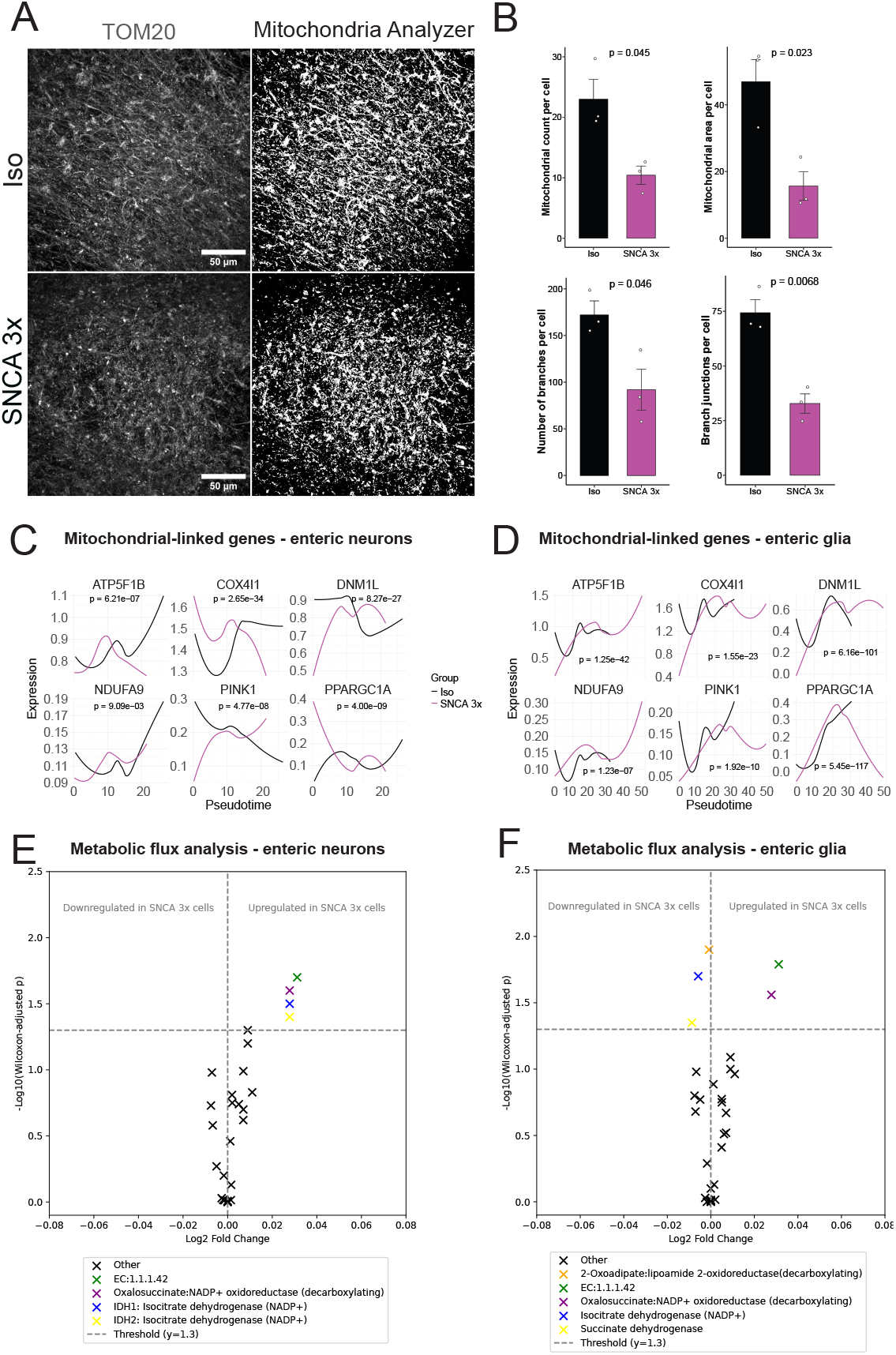
SNCA 3x drives basal mitochondrial dysfunction and alters enteric neuron-glia communication in iPSC-ENLs. **(A)** Representative panels of TOM20 immunostaining (left) and the mitochondrial mask created with the Mitochondria Analyzer plugin for FIJI. Uper panels represent Iso and bottom panels SNCA 3x ENLs. Cells were analyzed on day 70 after start of differentiation. **(B)** Quantification of mitochondrial count, area, number of branches and branch junctions in iPSC-ENLs calculated with the Mitochondria Analyzer plugin for FIJI. All data were normalized to the number of cell nuclei per image, calculated using DAPI staining in CellProfiler. At least 3 individual images per line were averaged, being the data representative of n=3 biological replicates per group, mean ± SEM, p-values calculated by unpaired two-tailed Student’s t-test. **(C and D)** Expression of mitochondrial-linked genes in SNCA 3x vs. Iso enteric neurons (C) and glia (D) along pseudotime. Gene expression values were modeled using linear regression across pseudotime, with FDR correction for multiple comparisons. **(E and F)** Metabolic flux analysis of mitochondrial function in SNCA 3x vs Iso enteric neurons (E) and glia (F). We calculated the logarithm of the Wilcoxon adjusted p-value to filter reaction differences. A threshold of 1.3 (corresponding to an adjusted p-value of 0.05) was used to identify significant reactions.

Using the scRNAseq data, we built on the previous analyzes and explored how SNCA 3x affects cellular communication and signaling in iPSC-ENLs using CellChat^45^ (Figures S1F-H, Figures S3C-H, Table S6). In enteric neurons, SNCA 3x reduced pathway interactions, particularly those linked to mitochondrial protection and energy metabolism, such as EGF^46^, and shifted signaling toward NOTCH^47^, which is linked to mitochondrial stress and inflammation (Figures S3C-E, Table S6). In contrast, SNCA 3x glial cells showed increased associations with pathways involved in mitochondrial repair and energy production, including EGF (Figures S3F-H, Table S6). This suggests that SNCA 3x enteric glial cells compensate for neuronal stress by enhancing mitochondrial support. Pathways like GDF^48^, CypA^49^, and WNT^50^ signaling further emphasize mitochondrial dysfunction in SNCA 3x neurons and the glial response to buffer this dysfunction through energy production and repair pathways (Figures S3C-H, Table S6). The findings highlight the balance between neuroprotective and neurotoxic mechanisms driven by increased α-syn, with glial cells playing a key compensatory role.

Together, these results establish basal mitochondrial dysfunction as a central feature of α-syn pathology in the ENS, characterized by structural, transcriptional, and metabolic alterations that reshape enteric neuron-glia interactions. These changes set the stage for further metabolic and functional impairments under inflammatory conditions, as explored in the following section.

### TNF-α uncovers genotype-specific α-syn accumulation and synaptic dysfunction in SNCA 3x ENLs

The ENS, as a vital component of the intestinal microenvironment, plays a role in coordinating local immune responses^51^, besides other functions. Cytokines, as key regulators of immune signaling, mediate both physiological and pathological processes within the ENS^51^.

Our scRNAseq data revealed that SNCA 3x alters biological pathways and intercellular communication. Additional ligand-receptor pair analysis identified activated inflammatory pairs (e.g., edn1-ednrb^52^, crlf1-lifr^53^) and repressed synapse-regulating pairs (e.g., nlgn1-nrxn3^54^, nxph1-nrxn1^55^) (Figure 4A) in SNCA 3x ENLs. This prompted us to investigate the impact of various cytokines (TNF-α, IL-10, IL-23, IL-17) – relevant both to PD and regulation of mucosal immunity^1,51^ - on α-syn biology and neuronal function in iPSC -ENLs. Cytokine treatments revealed a genotype-dependent effect, as only TNF-α significantly increased total α-syn levels in SNCA 3x ENLs compared to TNF-α-treated Iso controls, without affecting cell viability (Figures 4B-D). Dot blot analysis further confirmed higher α-syn levels in TNF-α-treated SNCA 3x ENLs compared to TNF-α-treated Iso ENLs (Figure S4A). However, there was no significant difference between SNCA 3x ENLs at baseline and TNF-α-treated SNCA 3x ENLs, indicating that the genotype itself determines α-syn elevation, with TNF-α further exacerbating this difference relative to controls. Given previous reports of α-syn aggregates in the gut of PD patients^8^, we assessed its solubility using sarkosyl fractionation^56^. α-syn signal was detected in the pellet fractions of SNCA 3x ENLs, but no significant differences were observed between Iso and SNCA 3x under basal conditions or following TNF-α stimulation (Figure S4B). We investigated further potential changes using ELISA, which revealed a modest increase in aggregated α-syn in SNCA 3x ENLs upon TNF-α stimulation (Figure 4E). Cytokine array showed no significant changes in cytokine production between groups upon TNF-α treatment

**Figure 4.**
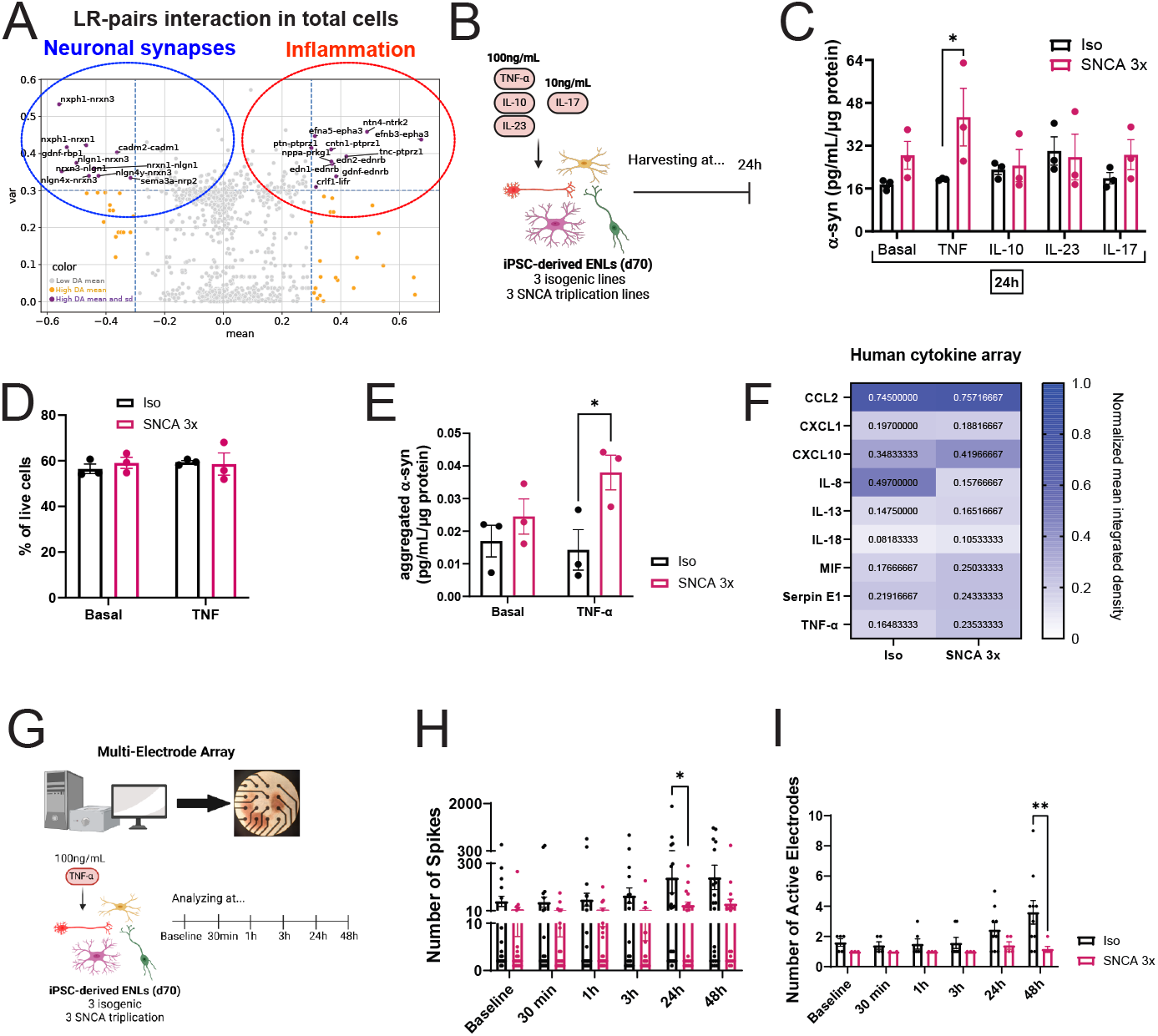
TNF-α uncovers genotype-specific α-syn accumulation and synaptic dysfunction in SNCA 3x ENLs. **(A)** Ligand-Receptor pair analysis of the scRNAseq data prior to subclustering. This plot depicts pairs that show both a large mean change as well as a large variance in the population (purple). Cells that don’t make the variance cut-off of 0.3 are shown in orange. Cells that don’t make the mean cut-off of 0.3 are shown in grey. **(B)** Experimental paradigm showing the stimulation of iPSC-ENLs with different cytokines. Generated using BioRender. **(C)** ELISA of total α-syn (Abcam, cat:ab260052) measured intracellularly in iPSC-ENLs, normalized by total protein. n=3 biological replicates per group, mean ± SEM, *p<0.05 by two-way ANOVA with Sidak post-hoc. Basal refers to cells treated with vehicle used to dilute the cytokines (DPBS+0.1% BSA). **(D)** Flow cytometry analysis of the percentage of live cells using Live/Dead staining. n=3 biological replicates per group, mean ± SEM. Basal refers to cells treated with vehicle used to dilute the TNF-α (DPBS+0.1% BSA). **(E)** ELISA of aggregated α-syn (Biolegend, cat:448807) quantified intracellularly in iPSC-ENLs, normalized by total protein. n=3 biological replicates per group, mean ± SEM, *p<0.05 by two-way ANOVA with Sidak post-hoc. Basal refers to cells treated with vehicle used to dilute the TNF-α (DPBS+0.1% BSA). **(F)** Human cytokine array quantification of the supernatants of iPSC-ENLs previously stimulated for 24h with 100 ng/ml TNF-α. Data represent the mean of n=3 biological replicates per group. **(G)** Experimental paradigm for the MEA experiments. Generated using BioRender. **(H and I)** Quantification of the number of spikes (H) and number of active electrodes (I) generated from the MEA data, n=wells of a CytoView MEA 48-well plate, representative of 3 biological replicates per group, mean ± SEM, *p<0.05, **p<0.01 by two-way ANOVA with Sidak post-hoc.

(Figure 4F), which were also not detected at the gene expression level (data not shown). This suggests that SNCA 3x ENLs exhibit an intrinsic susceptibility to TNF-α, with selective α-syn increase occurring independently of the induction of a broad inflammatory response.

Finally, to assess how TNF-α impacts enteric neuronal function over the time, we performed a functional electrophysiology assay to measure neuronal activity using the Multielectrode array (MEA) platform (Figure 4G). Interestingly, while the Iso ENLs reacted to TNF-α treatment by increasing the number of spikes and active electrodes over time, SNCA 3x ENLs showed no response (Figures 4H-I). There was a significant difference between groups after 24h and 48h of TNF-α exposure in number of spikes (Figure 4H) and active electrodes (Figure 4I), respectively. No differences were observed between groups regarding weighted mean firing rate measurements (Figure S4C).

These findings suggest that SNCA 3x ENLs exhibit an impaired capacity to modulate their neuronal activity in response to TNF-α. This functional deficit, coupled with selective α-syn increase, further supports an intrinsic vulnerability of SNCA 3x ENLs to inflammatory stress, reinforcing a link between α-syn accumulation and synaptic dysfunction in the ENS.

### TNF-α rewires the metabolism of SNCA 3x ENLs by disrupting the malate-aspartate shuttle and impairing TCA cycle flux

Building on the links between α-syn and mitochondrial dysfunction identified in our scRNAseq data, we used multiomics analyses to examine the impact of TNF-α on the iPSC-ENL proteome and metabolome (Figure 5A).

**Figure 5.**
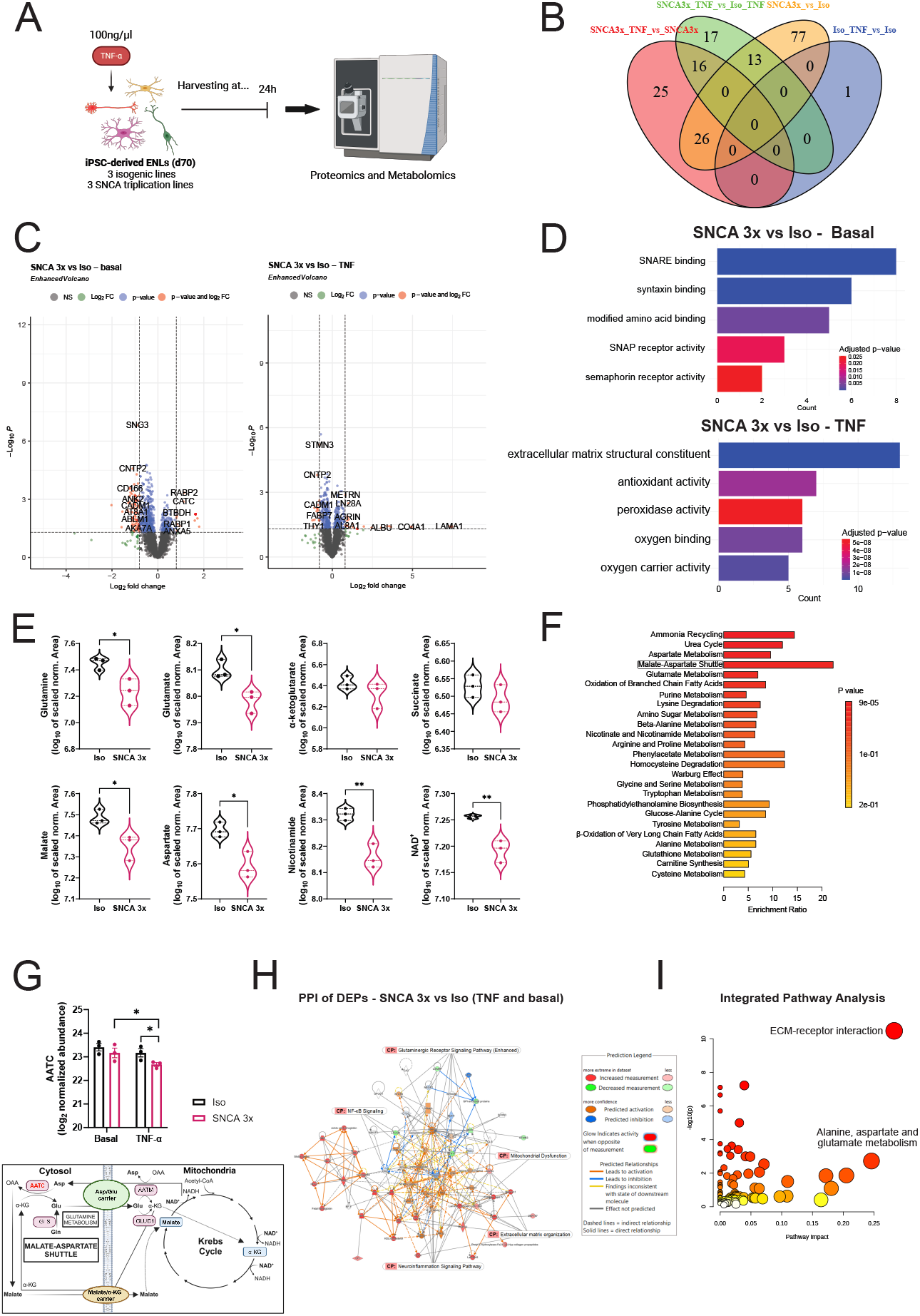
TNF-α rewires the metabolism of SNCA 3x ENLs by disrupting the malate-aspartate shuttle and impairing TCA cycle flux. **(A)** Experimental paradigm of the proteomics and metabolomics experiments. Proteomics data included both basal and TNF-α-stimulated iPSC-ENLs whereas metabolomics data consisted only of TNF-α-stimulated cells. Generated using BioRender. **(B)** Venn diagram showing the differentially expressed proteins (FC>1.75 and p<0.05) between 4 different comparisons: SNCA 3x vs Iso in a basal level (yellow), SNCA 3x vs Iso treated with TNF-α (green), SNCA 3x treated vs basal (red), Iso treated vs basal (blue). p-values were calculated using a linear model and empirical Bayes moderation (limma). See table S7. Basal refers to cells treated with vehicle used to dilute the TNF-α (DPBS+0.1% BSA). **(C)** Volcano plots highlighting proteins obtained as in (B) for two comparisons: SNCA 3x vs Iso in a basal level and SNCA 3x vs Iso treated with TNF-α. Basal refers to cells treated with vehicle used to dilute the TNF-α (DPBS+0.1% BSA). **(D)** Molecular function enrichment analysis of the comparisons highlighted in (C). See table S8. Basal refers to cells treated with vehicle used to dilute the TNF-α (DPBS+0.1% BSA). **(E)** Quantification of intracellular metabolites in iPSC-ENLs treated with TNF-α. p-values were calculated by performing a scaling of the data to consider the donor effect. n=3 biological replicates per group, mean ± SEM, *p<0.05, **p<0.01 by unpaired two-tailed Student’s t-test. **(F)** Metabolite set enrichment analysis on annotated compounds found by metabolomics analysis. See table S9. **(G)** Quantification of cytosolic aspartate aminotransferase (AATC) from the proteomics data. n=3 biological replicates per group, mean ± SEM, *p<0.05. p-values were calculated using a linear model and empirical Bayes moderation (limma). Basal refers to cells treated with vehicle used to dilute the TNF-α (DPBS+0.1% BSA). Below the graph is a metabolic map for an integrated visualization of the MAS, TCA cycle and glutamine metabolism. OAA=oxaloacetate, Asp=aspartate, Gln=glutamine, Glu=glutamate, αKG= α-ketoglutarate, GLS=glutaminase, GLUD1=glutamate dehydrogenase, AATM=mitochondrial aspartate aminotransferase. AATC is highlighted in red. **(H)** Protein-protein interaction network generated used Ingenuity Pathway Analysis by merging the networks of the following comparisons: SNCA 3x vs Iso –TNF-α treated vs basal. Colored nodes highlight the TNF-α induced effects. **(I)** Integrated pathway analysis generated by merging the differentially expressed proteins and metabolites from the comparison SNCA 3x vs Iso treated with TNF-α.

Data independent acquisition (DIA)-based shotgun proteomics coupled to label-free quantification was used to quantify the proteomes of Iso and SNCA 3x ENLs in a basal level and upon TNF-α stimulation. When these proteomes were compared (Figures 5B-C and Table S7), differentially expressed proteins (DEPs) were identified. The effect of SNCA 3x (Iso versus SNCA 3x, basal) resulted in the most robust proteomic alteration with 116 DEPs. This indicates a strong effect of α-syn in the proteome of iPSCs cells. Interestingly, only one DEP was identified when Iso cells were treated with TNF-α while 67 DEPs were identified in SNCA 3x cells upon TNF-α treatment (Table S7). This strongly suggests that α-syn sensitizes iPSCs-ENLs to cytokines, in line with the strong contribution of α-syn in SNCA 3x cells. Gene Ontology Molecular Function analyses revealed a significant enrichment in synaptic functions, including SNAP receptor and SNARE binding in basal SNCA 3x ENLs, suggesting neurotransmitter release and vesicle trafficking alterations^57^ (Figure 5D, Table S8). TNF-α exposure shifted the profile toward oxidative stress and mitochondrial dysfunction, marked by increased peroxidase, antioxidant, and oxygen transport functions, indicating impaired mitochondrial respiration (Figure 5D, Table S8). This progression from synaptic dysfunction to metabolic disruption underscores the impact of inflammatory stress.

LC-Orbitrap-MS metabolomics revealed reductions in aspartate, glutamate, malate and glutamine levels in SNCA 3x ENLs treated with TNF-α, suggesting malate-aspartate shuttle (MAS) impairment, a key process for tricarboxylic acid (TCA) cycle flux and mitochondrial redox balance^58^ (Figure 5E, Table S9). Accordingly, nicotinamide and NAD^+^ levels were significantly reduced in these cells, with no changes in NADH levels, suggesting altered NAD^+^ homeostasis, potentially due to impaired regeneration or increased utilization in response to metabolic stress (Figure 5E, Table S9). Metabolite-set enrichment analysis on the differentially expressed metabolites (DEMs) (Table S9) confirmed MAS as a top affected pathway (Figure 5F), supporting inflammation-driven metabolic reprogramming.

Proteomics further identified downregulation of aspartate aminotransferase (AATC) in TNF-α-treated SNCA 3x ENLs (Figure 5G), reinforcing MAS suppression and disrupted cytosol-mitochondria metabolic exchange. Protein-protein interaction (PPI) analysis interconnected neuroinflammatory pathways, including NF-κB activation and glutaminergic receptor signaling with mitochondrial dysfunction (Figure 5H). Integrated pathway analysis linked TNF-α exposure to extracellular matrix receptor interactions and glutamate metabolism (Figure 5I).

These findings demonstrate that TNF-α disrupts metabolic homeostasis in SNCA 3x ENLs by impairing MAS and the TCA cycle, leading to mitochondrial dysfunction.

### TNF-α increases α-syn-mitochondria interactions and induces oxidative stress in SNCA 3x ENLs

Given that α-syn is known to bind to mitochondria and that mitochondrial dysfunction can promote protein aggregation and oxidative stress^59^, we next investigated whether these metabolic alterations observed at a basal level (Figure 3) and with TNF-α treatment (Figure 5) contribute to mitochondrial α-syn accumulation and oxidative stress using several techniques (Figure 6A). Proximity Ligation Assay (PLA) of α-syn and TOM20 confirmed increased α-syn-mitochondria interactions in SNCA 3x ENLs, which were further amplified by TNF-α stimulation (Figures 6B-C). Next, we successfully isolated the mitochondria of iPSC-ENLs, as indicated by a strong and uniform TOM20 immunostaining across all samples (Figure 6D). Subsequently, we used ELISA to measure the total α-syn levels in the mitochondrial and cytosolic fractions. We observed that even at basal conditions, α-syn levels are significantly increased in the mitochondria of SNCA 3x ENLs. The same trend was observed following TNF-α stimulation, although not significant (Figure 6E). Finally, to investigate whether TNF-α could also increase mitochondrial stress, we analyzed superoxide production using flow cytometry of MitoSox. We observed that upon stimulation, superoxide levels were significantly higher in SNCA 3x ENLs (Figure 6F).

**Figure 6.**
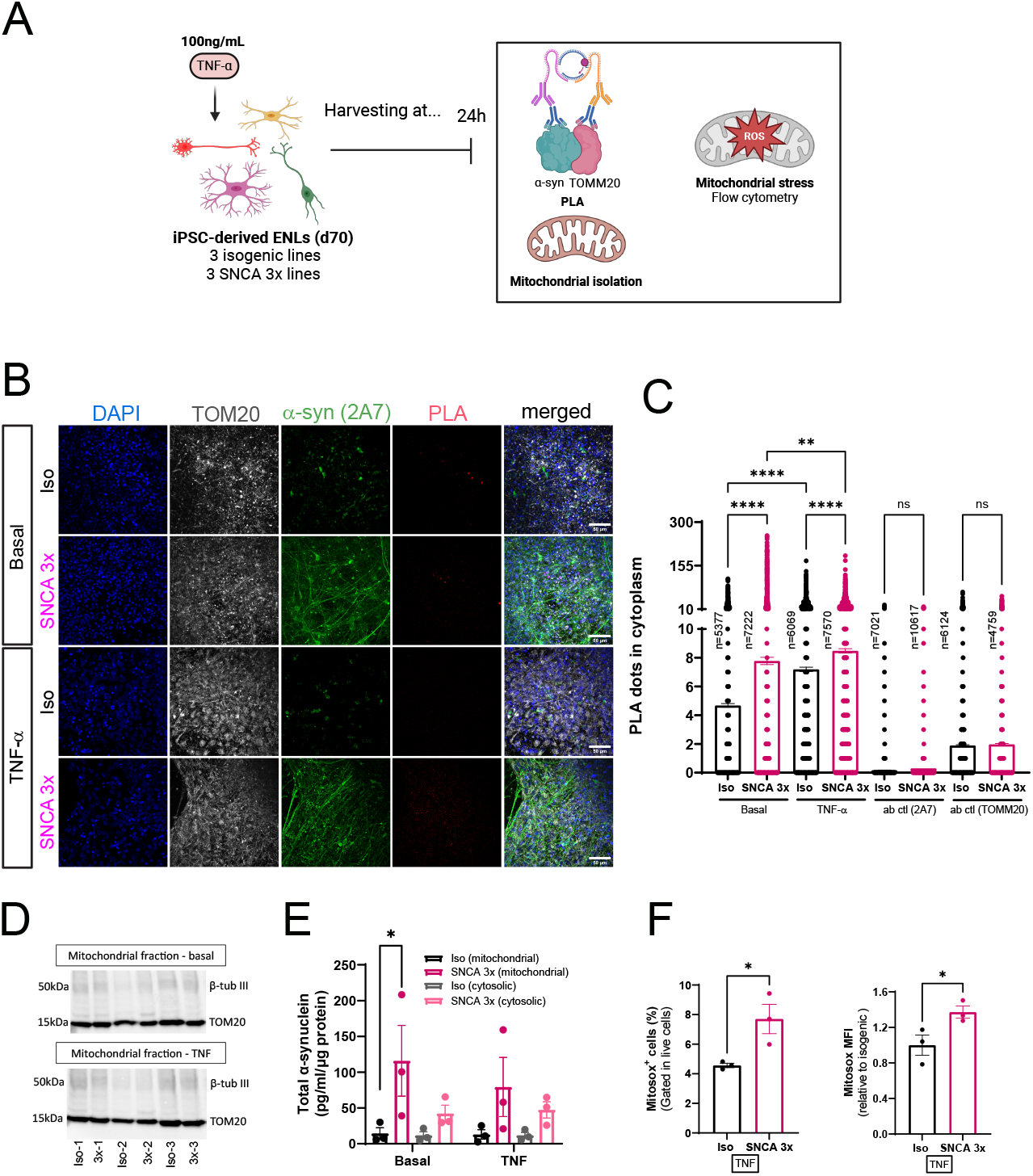
TNF-α increases α-syn-mitochondria interactions and induces oxidative stress in SNCA 3x ENLs. **(A)** Paradigm describing the PLA, mitochondrial isolation and measurement of mitochondrial ROS experiments. Generated using BioRender. **(B)** Assessment of PLA results using immunocytochemistry. Panels are separated in basal and TNF-α stimulated for both Iso and SNCA 3x groups. Panels show the nuclear marker DAPI (blue), the mitochondrial marker TOM20 (gray), α-syn (green) and PLA dots for TOM20-α-syn (red). Basal refers to cells treated with vehicle used to dilute the TNF-α (DPBS+0.1% BSA). **(C)** Quantification of the PLA dots in the cytoplasm. n= 3 biological replicates per group, with number of total individual cells analyzed per condition indicated in the figure, mean ± SEM, **p<0.01, ****p<0.0001, by one-way ANOVA with Tukey’s post-hoc. Basal refers to cells treated with vehicle used to dilute the TNF-α (DPBS+0.1% BSA). **(D)**Western-blotting analysis for characterization of the mitochondrial isolation protocol using the beta-tubulin III and TOM20 markers for cytosol and mitochondria, respectively. Basal refers to cells treated with vehicle used to dilute the TNF-α (DPBS+0.1% BSA). **(E)** ELISA of total α-syn quantified in the mitochondrial isolates of iPSC-ENLs, normalized by total protein. n=3 biological replicates per group, mean ± SEM, *p<0.05 by two-way ANOVA with Sidak post-hoc. Basal refers to cells treated with vehicle used to dilute the TNF-α (DPBS+0.1% BSA). **(F)** Flow cytometry analysis of the percentage and median fluorescence intensity (MFI) of superoxide producing cells in TNF-α treated ENLs using MitoSox staining. n=3 biological replicates per group, mean ± SEM, *p<0.05 by unpaired two-tailed Student’s t-test

These findings establish a direct link between TNF-α-induced metabolic reprogramming and mitochondrial pathology, where increased oxidative stress and enhanced α-synmitochondria interactions may contribute to impaired mitochondrial function and bioenergetic deficits, further worsening neuronal dysfunction in SNCA 3x ENLs.

### TNF-α increases the reliance of SNCA 3x ENLs on glutamine oxidation, which is rescued by CSB6 treatment

Next, we investigated α-syn-mitochondria interactions in iPSC-ENLs by assessing mitochondrial function following TNF-α stimulation using Seahorse and flow cytometry assays. We also tested the potential of the glutamate metabolism modulator, CSB6, to restore mitochondrial function in our model (Figure 7A).

**Figure 7.**
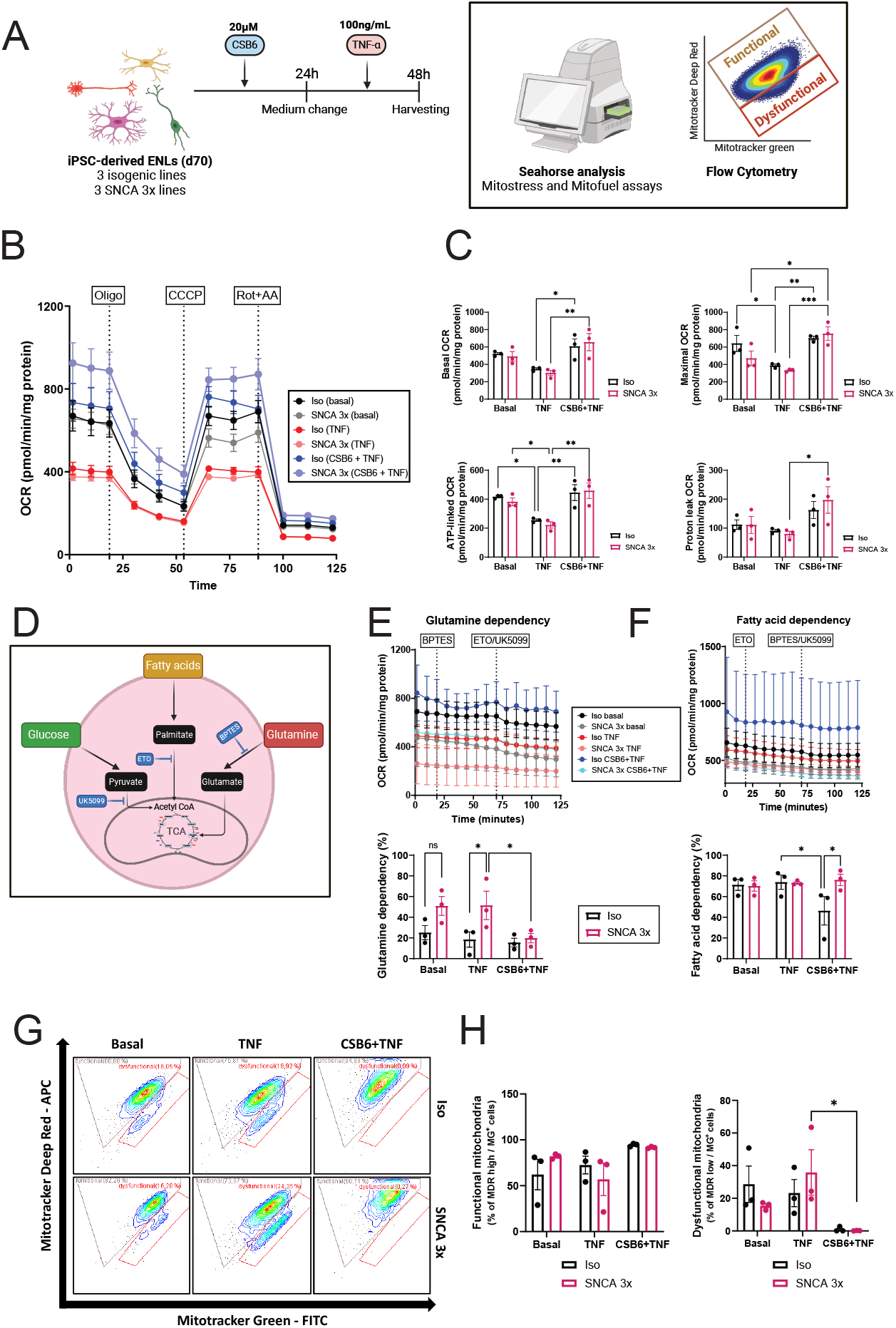
TNF-α increases the reliance of SNCA 3x ENLs on glutamine oxidation, which is rescued by CSB6 treatment. **(A)** Paradigm of the functional experiments using seahorse and mitotracker assays. iPSC-ENLs were treated with CSB6 for 24h, then the medium was replaced by one containing TNF-α for further 24h. Generated using BioRender. **(B)** Mitostress assay. Curves showing the overall OCR for Iso and SNCA 3x ENLs with the different treatments specified. Oligo: Oligomycin; CCCP: carbonyl cyanide m-chlorophenylhydrazone; AA+Rot – antimycin-A plus Rotenone. Basal refers to cells treated with vehicle used to dilute the TNF-α and CSB6 (DPBS+0.1% BSA). **(C)** Quantification of basal, maximal, ATP-linked and proton leak OCR from the measurements seen in (B). Data was normalized to total protein. n=3 biological replicates per group, mean ± SEM, *p<0.05, **p<0.01, ***p<0.001, by two-way ANOVA with Tukey’s post-hoc. Dots represent the average for each cell line of three independent experiments. Basal refers to cells treated with vehicle used to dilute the TNF-α and CSB6 (DPBS+0.1% BSA). **(D)**Experimental paradigm illustrating the Seahorse mitofuel assay. BPTES= Bis-2-(5-phenylacetamido-1,3,4-thiadiazol-2-yl)ethyl sulfide, ETO=etomoxir. Generated using BioRender. **(E and F)** OCR curves and quantification of the glutamine (E) and fatty acid (F) dependency analysis. n=3 biological replicates per group, mean ± SEM, *p<0.05 by two-way ANOVA with Sidak post-hoc. Basal refers to cells treated with vehicle used to dilute the TNF-α and CSB6 (DPBS+0.1% BSA). **(G and H)** Flow cytometry analysis of mitochondrial function using Mitotracker probes in Iso and SNCA 3x ENLs with the different treatments specified (G) and quantification of functional and dysfunctional mitochondria (H). n=3 biological replicates per group, mean ± SEM, *p<0.05 by two-way ANOVA with Tukey’s post-hoc. Basal refers to cells treated with vehicle used to dilute the TNF-α and CSB6 (DPBS+0.1% BSA).

We measured mitochondrial oxidative phosphorylation (OX-PHOS) using the Seahorse mitostress assay: TNF-α-treated SNCA 3x ENLs exhibited reduced mitochondrial respiration, particularly in ATP-linked oxygen consumption rate (OCR), linking TNF-α to impaired mitochondrial metabolism and energy production (Figures 7B-C). CSB6 treatment fully restored mitochondrial capacity, significantly increasing basal, maximal, ATP-linked, and proton-leak OCR beyond baseline levels (Figures 7B-C). Notably, CSB6 selectively induced proton-leak OCR in SNCA 3x ENLs, suggesting a protective role via enhanced electron transport chain efficiency.

To determine metabolic substrate dependency, we performed a Seahorse Mitofuel assay, an analytical method for measuring mitochondrial fuel oxidation in live cells. Here, the oxidation of glucose, glutamine, and fatty acids is measured independently through the use of specific inhibitors (Figure 7D). TNF-α-treated SNCA 3x ENLs showed a significant increase in glutamine oxidation dependency (Figure 7E), consistent with the observed MAS impairment (Figure 5). CSB6 treatment not only rescued this TNF-α-induced glutamine reliance (Figure 7E), but also promoted a metabolic shift toward fatty acid oxidation (Figure 7F), suggesting enhanced mitochondrial substrate flexibility.

Lastly, to assess functional mitochondrial integrity, we performed flow cytometry analysis using MitoTracker dyes (Figures 7G-H). TNF-α-treated SNCA 3x ENLs displayed a trend toward increased dysfunctional mitochondria (MitoGreen^+^/MitoDeepRed^low^), suggesting a loss of mitochondrial membrane potential. Importantly, CSB6 treatment rescued this phenotype, significantly decreasing the dysfunctional mitochondrial population and restoring mitochondrial health in SNCA 3x ENLs (Figures 7G-H).

In summary, our findings demonstrate that TNF-α reprograms metabolism in SNCA 3x ENLs by impairing MAS, reducing TCA cycle flux, and shifting mitochondrial dependency toward glutamine oxidation as a compensatory mechanism. This metabolic stress is linked to increased mitochondrial α-syn accumulation, oxidative stress, and loss of mitochondrial membrane potential. CSB6 counteracts these TNF-α-driven alterations by restoring mitochondrial respiration, reducing glutamine dependency, enhancing substrate flexibility, and preventing mitochondrial dysfunction. The results identify a TNF-α-induced metabolic vulnerability in SNCA 3x ENLs and highlight CSB6 as a potential therapeutic intervention targeting mitochondrial function.

## Discussion

In this study, we employed a multiomics approach integrating single-cell transcriptomics, proteomics, metabolomics, and functional assays to gain a comprehensive understanding of α-syn-induced dysfunction in the human ENS, something that has been challenging to achieve with previous methodologies^60,61^. Our findings reveal that α-syn increases susceptibility of iPSC-derived ENLs to TNF-α, leading to mitochondrial dysfunction and metabolic reprogramming, characterized by a disruption in MAS and the TCA cycle, forcing ENLs to rely on glutamine oxidation for energy production. These data demonstrate that α-syn accumulation in ENLs drives vulnerability in the ENS through this inflammatory-metabolic crosstalk. Notably, targeting glutamate metabolism with CSB6 restored mitochondrial metabolism, highlighting a potential therapeutic approach for early PD-related ENS dysfunction.

Our scRNAseq analysis revealed that SNCA 3x induces mitochondrial dysfunction at the transcriptional level, affecting both enteric neuron and glia subpopulations. Neuronal subclusters exhibited significant downregulation of *ATP5F1B* and *COX4I1*, genes essential for ATP synthase function and cytochrome c oxidase activity^40^, respectively, suggesting compromised OXPHOS capacity. In parallel, upregulation of *PINK1* and *DNM1L* indicates a shift toward mitophagy^43^ and mitochondrial fission^44^, processes associated with mitochondrial network fragmentation under stress. The TOM20 analysis further supports these findings, showing a reduction in mitochondrial count and area, reinforcing that SNCA 3x ENLs undergo structural mitochondrial remodeling in response to α-syn-induced metabolic stress^62^. In contrast, enteric glial cells showed a transient upregulation of mitochondrial biogenesis genes, alongside enhanced levels of OXPHOS-linked genes (*ATP5F1B, COX4I1, NDUFA9*). These alterations potentially provide metabolic support to stressed neurons, indicating a compensatory response aimed at buffering mitochondrial dysfunction. Together, these findings demonstrate that at basal levels, α-syn accumulation induces metabolic disruptions in enteric neurons, whereas enteric glia activate compensatory mechanisms to support neuronal function, recapitulating the previously described increase in neuron-glia crosstalk observed as a response to α-syn toxicity^63^. In line with this, most of our enteric glial cells corresponded to the Glia1 subtype described by Drokhlyansky et al.^26^, associated with GDNF signaling, which has been suggested as a compensatory mechanism in early PD pathology^64^.

In our iPSC-ENL model, we detected that TNF-α unmasked genotype-specific α-syn accumulation in SNCA 3x ENLs, despite not significantly changing the immunoreactivity of the cells. Interestingly, differences in α-syn aggregation were only detectable using highly sensitive ELISA assays, suggesting that our model represents a pre-aggregation stage. In contrast to classical CNS models^24^, the presence of enteric glial cells in ENLs suggests the activation of compensatory mechanisms that may help buffer excessive α-syn aggregate formation. Additionally, α-syn was previously described primarily in its monomeric form in the ENS, indicating that the native state of the protein differs from what is observed in the CNS^65^. Notably, TNF-α levels are elevated in the serum of PD patients and correlate with disease progression^66,67^. Moreover, TNF-α is synthesized by intestinal epithelial cells^68^, suggesting that it can locally impact gut inflammation and enhance α-syn accumulation in the ENS. A compelling link between intestinal TNF-α and PD pathophysiology comes from studies showing that treatment with gold-standard anti-TNF therapies in IBD patients reduced PD risk by nearly 80%^69^. Although TNF-α did not exert major effects in terms of neuronal activity, our MEA data suggested a strong basal synaptic dysfunction phenotype in SNCA 3x ENLs, an important observation given that over 80% of PD patients experience constipation, which can be directly associated with ENS activity dysfunction^70^.

Our multiomics analysis not only validated the scRNAseq findings but also provided deeper insights into a TNF-α-induced metabolic rewiring in SNCA 3x ENLs, bringing important novel insights into metabolic vulnerability in the ENS in the context of early PD. Using proteomics, we observed that increased α-syn impacted neuronal function, in line with the previously mentioned transcriptomics signatures as well as the neuronal activity data. This is also in line with recent findings using cortical organoids^71^. The addition of TNF-α highlighted oxidative stress and mitochondrial dysfunction as a response to inflammation-driven α-syn toxicity. Additionally, using metabolomics, we uncovered a hypometabolic state in SNCA 3x ENLs, characterized by a major downregulation in metabolites related to TCA and MAS. In particular, we found reduced NAD^+^ levels, with NADH remaining unaltered, suggesting impaired NAD^+^ regeneration under inflammatory stress, as observed in PD and other neurodegenerative diseases^72-74^. Given that NAD^+^ is a critical cofactor for TCA cycle enzymes such as malate dehydrogenase and its role in electron transport chain function, its depletion could directly impair mitochondrial substrate oxidation and ATP production. The failure to replenish NAD^+^ pools impairs mitochondrial respiration, likely through disruptions in the MAS, which facilitates the transfer of cytosolic NADH into mitochondria to maintain redox balance and sustain OXPHOS. These findings align well with a previous study using iPSC-derived neuronal models of sporadic PD^75^, which also highlighted cellular metabolic disruption as a clear phenotype. Interestingly, in line with our proteomics data, expression of the *GOT1* gene, coding for AATC, was lower in the brain of PD patients compared to controls^76^. Also corroborating these results, a study using a mouse model for prodromal PD showed that TCA dysfunction influences the onset and progression of α-syn pathology^77^. Therefore, our findings emphasize the inflammatory-driven metabolic reprogramming in SNCA 3x ENLs, pointing to the interplay of inflammation, mitochondrial dysfunction, and MAS/TCA cycle disruption in enteric α-synucleinopathies.

Deepening into the mechanism, our data indicated that already at basal levels, there is an increased interaction between α-syn and mitochondria in SNCA 3x ENLs, in line with previous findings^78^, which is further enhanced by the addition of TNF-α in the system. Additionally, TNF-α increased mitochondrial superoxide production in SNCA 3x ENLs, reinforcing the role of inflammation in promoting mitochondrial dysfunction^79^. We also observed that TNF-α has a general pattern of disrupting mitochondrial function, although no significant differences were noticed between Iso and SNCA 3x groups. Nevertheless, more specific analyses allowed us to uncover a mechanistic effect of TNF-α on rewiring the metabolic profile of SNCA 3x ENLs towards glutamine oxidation, characterizing an activation of anaplerotic pathways that are used to fuel cellular metabolism under stress conditions^80^. This can be explained by the observed MAS dysfunction, which limits aspartate availability for transamination reactions, ultimately restricting metabolic flexibility and forcing SNCA 3x ENLs to rely on glutamine oxidation to sustain TCA cycle activity. Finally, using our scRNAseq as reference, we predicted the compound CSB6, a vesicular glutamate transporter inhibitor that showed protective effects in previous studies with PD models^81^, as a candidate to rescue the phenotypes observed in SNCA 3x ENLs using a data-driven approach. Not only did CSB6 rescue the TNF-α-driven dependency on glutamine oxidation in SNCA 3x ENLs, but it also promoted a metabolic shift towards fatty acid oxidation, a more efficient pathway in terms of mitochondrial bioenergetics^82^, and increased overall mitochondrial function.

In conclusion, our findings indicate that α-syn accumulation and TNF-α-induced inflammation synergize to drive ENS dysfunction, supporting a dual-hit hypothesis for PD onset. This interplay reinforces the ENS as an active player in PD pathophysiology, shifting the paradigm from a passive receptor of inflammatory signals to an initiator of metabolic disruption. Our iPSC-ENL model establishes a powerful platform for investigating early-stage PD mechanisms and suggests that targeting glutamate metabolism may offer a new therapeutic perspective for ENS-related dysfunction in PD.

### Limitations of the study

While the iPSC derived ENL broadly recapitulates the human ENS, the presence of mixed neuronal and glial cells in iPSC-ENLs makes it challenging to separate mechanistic effects in a populational level. We tried to address this by extensively analyzing our scRNAseq data. Additionally, it is very difficult to obtain gut specimen from early PD patients that fully capture the ENS extension. CSB6 is a compound that increased overall mitochondrial function in our model and there is evidence for effectiveness coming from other PD models^81^. Thinking about clinical translation, CSB6 served as a proof-of-concept in our study and previously approved compounds could be repurposed to counteract the metabolic alterations observed in the iPSC-ENLs in future approaches. Despite these limitations, our research provides a solid basis for comprehending enteric neural vulnerability in PD and highlights metabolic interventions as promising early therapeutic strategies.

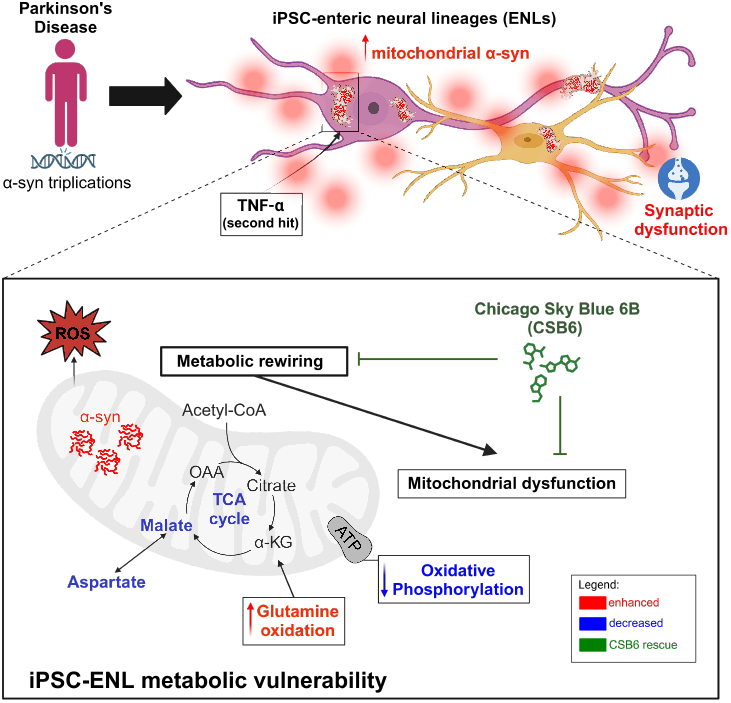

### Graphical summary

- SNCA 3x induces basal mitochondrial and synaptic dysfunction in iPSC-ENLs
- TNF-α exacerbates mitochondrial α-syn interactions, disrupting the malate-aspartate shuttle (MAS) and shifting metabolism toward glutamine oxidation
- Metabolic disruption in SNCA 3x ENLs is linked to oxidative stress and neuronal vulnerability under cytokine-induced stress
- CSB6 restores mitochondrial function and reverses TNF-α-driven metabolic rewiring in SNCA 3x ENLs

## Resource availability

Further information and requests for resources and reagents should be directed to and will be fulfilled by the lead contact, Beate Winner. All OMICs data generated from this study will be deposited and made public by the time of the final publication.

## Author contributions

B.G. designed the research, performed experiments, analyzed and interpreted the data, executed bioinformatics analyzes, created the figures, and wrote the manuscript. L.E.G., V.R., C.J., E.G, A.G., J.G. performed experiments and analyzed the data. H.W. and M.F. maintained iPSC cultures. M.K., M.S. and A.H. performed bioinformatic analysis on the scRNAseq data. F.Z. generated the iPSC lines used in this study in the laboratory of J. R. Mazzulli, Northwestern University, Chicago. T.R., I.P., J.W., D.B.B., C.G., F.J.T., R.R. and M.N. provided resources and supervised the project. P.G. wrote the manuscript, analyzed and interpreted data and supervised the bioinformatics analyzes. B.W. designed the research, provided resources, interpreted the data, wrote the manuscript, acquired funding and supervised the project. All authors critically reviewed and approved the final version of the manuscript.

## ACKNOWLEDGEMENTS

This work was Funded by the Deutsche Forschungsgemeinschaft (DFG, German Research Foundation) - CRU5024 (A01, A02, A04, B02) – Projektnummer 505539112. Further support came from the Bavarian Ministry of Science and the Arts in the framework of the ForInter network. The LC-MS system used for metabolomic analysis was funded by the Deutsche Forschungsgemeinschaft (DFG, German Research Foundation) – INST 90/1048-1 FUGG. A.H. and D.B.B. were supported by the German Federal Ministry of Education and Research (BMBF) – 031L0309A. M.S. and F.J.T.: Funded by the European Union (ERC, DeepCell - 101054957).

## Methodology

### iPSC generation, characterization and culture methods

#### iPSC reprogramming, generation of isogenic controls

B-lymphocytes containing a triplication in the *SNCA* gene were obtained from the Coriell NINDS and NIGMS Human Genetic Cell Repositories: GM15010 (3x-1), ND00196 (3x-2), ND00139 (3x-4, referred as 3x-3 throughout this paper). Further phenotypic and genotypic details on these lines and subjects are available on https://www.coriell.org under the beforementioned accession numbers. Reprogramming into iPSC was performed by transfection of B-lymphocytes with episomal plasmids containing Oct3/4, L-Myc, Sox2 and Klf4, as previously described^24^. Isogenic controls were generated using a dual nickase CRISPR/Cas9 strategy to disrupt exon 2 of the *SNCA* gene, fully characterized and validated in a previous publication^24^. Copy number variation and cell karyotyping analyzes revealed absence of chromosomal abnormalities. Iso and SNCA 3x iPSC lines were generated in the laboratory of Joseph R. Mazzulli (The Ken and Ruth Davee Department of Neurology, Northwestern University Feinberg School of Medicine, Chicago, IL 60611, USA) and provided to this study by Friederike Zunke.

#### iPSC culture maintenance

iPSCs were cultured in Geltrex (Thermo Fisher)-coated 6-well plates with Essential 8 Flex medium (Thermo Fisher), maintained at 37°C and 5% CO2, with media change performed every two days. For thawing, cryovials were warmed in a 37°C water bath, and cells were resuspended in fresh medium containing 10µM ROCK inhibitor (Y-27632 dihydrochloride, Tocris). Passaging occurred at 80%-90% confluency using ReLeSR (Stemcell technologies), with cells split at a 1:2 or 1:3 ratio. Cells were routinely tested for mycoplasma infection. For freezing, cells were detached with ReLeSR and resuspended in a freezing medium containing knockout serum replacement (KOSR, Thermo Fisher) + 10% DMSO before storage at –150°C.

#### iPSC characterization

iPSCs were characterized using flow cytometry staining for Nanog and Lin28A. iPSCs were initially treated with Accutase (Gibco, USA), washed with 1× PBS, centrifuged at 300g at room temperature for 5min and then resuspended in FC buffer (2% FCS, 0.01% sodium azide in PBS). We then counted 500,000 cells per line and proceeded with fixation and permeabilization with 100µl BD Fixation/Permeabilization Solution (BD Biosciences) for 10 minutes, followed by addition of 1ml BD Perm/Wash Buffer (BD Biosciences), incubation for 5 minutes and centrifugation at 300g for 3min. For intracellular staining of iPSCs, we used a combination of Nanog (cat: 130-120-704, Miltenyi) and Lin28A (cat:563597, BD Biosciences) primary antibodies (both 1:100) in BD Perm/Wash Buffer for 30 minutes, followed by washing and resuspension in 300µl FACS buffer. Cells were then acquired using a Cytoflex S cytometer (Beck-man Coulter) and analyzed with the CytExpert 2.4 software. At least 30,000 events per sample were analyzed.

### Differentiation of iPSCs into ENLs

The ENS differentiation protocol was adapted from Barber et al.^22^, using the novel chemically defined method. ENS induction began on day 0 with a nearly confluent iPSC monolayer and continued for 12 days with stepwise culture transitions. On day 0, Essential 8 medium (Thermo Fisher) was replaced with 2 ml of Cocktail A [Essential 6 medium (Thermo Fisher), 10µM SB431542 (Miltenyi), 600nM CHIR99021 (Tocris), 1ng/ml BMP4 (Peprotech), 100µg/mL normocin (InvivoGen)], followed by a switch to Cocktail B [Essential 6 medium (Thermo Fisher), 10 µM SB431542, 1.5µM CHIR99021, 100µg/mL normocin] on day 2, with media renewal on day 4. On day 6, Cocktail B was replaced with 4ml of Cocktail C [Essential 6 medium (Thermo Fisher), 10µM SB431542, 1.5µM CHIR99021, 1µM retinoic acid (Sigma), 100µg/mL normocin], with subsequent exchanges on days 8 and 10. On day 12, ENC spheroids were formed by detaching the ENC monolayer using Accutase for 30 min at 37°C, followed by centrifugation at 300g for 3 min, resuspension in NC-C medium [Neurobasal medium (Thermo Fisher), 1x N2 supplement (Thermo Fisher), 1x B27 supplement w/o Vit. A (Thermo Fisher), 1x GlutaMax (Thermo Fisher), 1x MEM non-essential aminoacids (Thermo Fisher), 10 ng/ml FGF2 (Peprotech), 3 µM CHIR99021, 100µg/mL normocin], and transfer to ultra-low attachment (ULA) plates. Spheroids formed within two days, with media replacement using NC-C medium. On day 15, EN induction began, involving dissociation of spheroids using Accutase, centrifugation at 300g, and resuspension in EN-C medium [Neurobasal medium (Thermo Fisher), 1x N2 supplement, 1x B27 supplement with Vit. A (Thermo Fisher), 1x GlutaMax, 1x MEM non-essential aminoacids, 100µM ascorbic acid (Sigma), 10ng/ml GDNF (Peprotech), 100µg/mL normocin]. Cells were counted using a fluorescence cell counter and seeded onto Geltrex-coated plates at 60,000 cells/cm^2^.

### RNA extraction, cDNA synthesis and RT-qPCR

Reverse transcription-quantitative PCR (RT-qPCR) analysis was performed starting with RNA extraction using Trizol (Invitrogen) and the RNeasy Mini Kit (Qiagen). Cells were homogenized in Trizol, followed by chloroform addition and phase separation by centrifugation at 12000g, 4°C for 15 min. The RNA-containing aqueous phase was collected, mixed with 70% ethanol, and processed through RNeasy columns with sequential washes using buffer RW1 and buffer RPE. RNA was eluted with RNase-free water, its concentration measured using the NanoPhotometer NP80 (Implen), and stored at -80°C. For cDNA synthesis, the QuantiTect Reverse Transcription Kit (Qiagen) was used, with 1000ng starting RNA concentration per reaction. After DNA removal with gDNA wipeout buffer, reverse transcription was carried out at 42°C for 30 min, followed by inactivation at 95°C for 3 min, and storage at -20°C. RT-qPCR was performed in triplicates for target genes (*TUBB3, GFAP, PHOX2B, ELAVL4, HOXB3*), with *HPRT* and *RPLP0* as housekeeping genes. Primer sequences used are listed in Table S1. A primer mix (10 µM) was prepared, and 1 µl cDNA per sample was loaded onto a 384-well plate. The qPCR master mix was added, the plate was sealed, centrifuged, and run on the LightCycler480 equipment (Roche). Ct values were analyzed using the delta-delta Ct method, and fold change values were calculated based on the expression values of the Iso group at day 6 of differentiation.

### Immunocytochemistry

Cells were fixed on day 70 after start of differentiation using 4% paraformaldehyde (PFA) for 30 min at room temperature (RT) (500 µl per well in an ibidi 24-well plate), washed three times in DPBS, and stored at 4°C until staining. Permeabilization was performed with 0.3% TritonX-100 (Sigma) in DPBS (Thermo Fisher) for 10 min at RT, followed by blocking with 5% donkey serum in DPBS with 0.3% TritonX-100 for 30 min. Cells were then incubated overnight at 4°C with the primary antibodies (mouse anti-a-syn, Novus, cat: NBP1-05194, 1:100; mouse anti-HuC/D, Thermo Fisher, cat: A-21271, 1:200 and rabbit anti-GFAP, DAKO, cat: Z0334, 1:500). The following day, cells were washed, incubated for 2 h at RT with secondary antibodies (AF488 anti-mouse; AF647 anti-rabbit, Thermo Fisher, all 1:500), and counterstained with DAPI (1:5000 in DPBS) for 2 min. After final washes, samples were mounted in Mowiol for long-term storage at 4°C. Imaging was performed using the Super Resolution Evident SR Spinning Disk microscope (Olympus) and the cellSens Software and processing was performed using FIJI (ImageJ).

### Single cell RNA sequencing

#### Sample preparation and sequencing

On day 70 of differentiation, we obtained single cells from iPSC-ENLs using a papain-DNAse digestion (cat: LK003150, Worthington) protocol. Cells were then cryopreserved in a medium containing 90% fetal bovine serum and 10% DMSO and sent out for library preparation and sequencing at GENEWIZ (Leipzig, Germany) with 6,000 cells targeted per sample and 50,000 reads targeted per cell. RNA samples were quantified using Qubit 4.0 Fluorometer (Life Technologies, Carlsbad, CA, USA) and RNA integrity was checked with RNA Kit on Agilent 5300 Fragment Analyzer (Agilent Technologies, Palo Alto, CA, USA).

rRNA depletion was performed using NEBNext rRNA Depletion Kit (Human/Mouse/Rat). RNA sequencing library preparation used NEBNext Ultra RNA Library Prep Kit for Illumina by following the manufacturer’s recommendations (NEB, Ipswich, MA, USA). Briefly, enriched RNAs were fragmented according to manufacturer’s instruction. First strand and second strand cDNA were subsequently synthesized. cDNA fragments were end repaired and adenylated at 3’ends, and universal adapter was ligated to cDNA fragments, followed by index addition and library enrichment with limited cycle PCR. Sequencing libraries were validated using NGS Kit on the Agilent 5300 Fragment Analyzer (Agilent Technologies, Palo Alto, CA, USA), and quantified by using Qubit 4.0 Fluorometer (Invitrogen, Carlsbad, CA).

The sequencing libraries were multiplexed and loaded on the flow cell on the Illumina NovaSeq X plus instrument according to manufacturer’s instructions. The samples were sequenced using a 2×150 Pair-End (PE) configuration v1.5. Image analysis and base calling were conducted by the NovaSeq Control Software v1.7 on the NovaSeq instrument. Raw sequence data (.bcl files) generated from Illumina NovaSeq was converted into fastq files and de-multiplexed using Illumina bcl2fastq program version 2.20. One mismatch was allowed for index sequence identification.

#### Initial processing, quality control, and clustering

The fastq files were processed using CellRanger (v7.1.0) from 10X Genomics, with reads aligned to the GRCh38 genome assembly to generate gene counts per cell. Subsequent processing, analysis, and visualization were performed in R (v4.3.3) using Seurat (v5.1.0). To ensure quality control, outlier cells with < 200 genes, > 7500 reads or a mitochondrial percentage > 5% were filtered out. Potential doublets were removed using *scDblfinder*. Following quality control, batch effect across samples was mitigated using Seurat’s integration workflow. Data normalization was carried out using the *SCTransform* algorithm. Features to use when integrating multiple datasets were identified using *SelectIntegrationFeatures*. Integration anchors computed using the *FindIntegrationAnchors* function were subsequently used by the *IntegrateData* function to generate a batch-corrected, integrated expression matrix. Principal component analysis (PCA) was applied for dimensionality reduction. Clustering was subsequently performed computing the shared nearest neighbor (SNN; *FindNeighbors*, k = 20) and applying the graph-based Leiden algorithm (*FindClusters*, resolution = 0.1) to determine the cell subtypes, followed by visualization with Uniform Manifold Approximation and Projection (UMAP). Initial annotation of the clusters was performed based on the expression of canonical markers^22,25^.

#### Subclustering analysis

To further resolve cellular heterogeneity, subclustering was performed on the enteric neurons and glia cell populations using Seurat (v5.1.0). Here, the enteric neurons and glial clusters were extracted from the integrated Seurat object using the subset function. Data normalization was conducted using the *NormalizeData* function, followed by identification of highly variable features via *FindVariableFeatures* with the ‘vst’ selection method. As in the previous analysis, the data was scaled, and PCA-based dimensionality reduction was performed. Clustering was carried out by computing the SNN (*FindNeighbors*, k = 20) and applying the Leiden algorithm (*FindClusters*, resolution 0.5). The processed objects were saved in H5Seurat format using the *SaveH5Seurat* function.

#### Mapping of scRNAseq dataset to a reference dataset

Mapping of the scRNAseq dataset to the reference scR-NAseq dataset from Drokhylansky et al.^26^ was performed using SingleR^83^ for cell-type annotation. The raw counts and the corresponding cell-type annotations from the reference dataset were converted to a SingleCellExperiment (SCE) object and log-normalized using *logNormCounts*. Query datasets of enteric neurons and glia were similarly transformed into SCE objects and subsetted into isogenic and mutant conditions. SingleR classification was conducted separately for Iso and SNCA 3x samples to obtain cell-type predictions. Visualization included heatmaps of classification scores (*plotScoreHeatmap*) and log-transformed assignment frequencies (*pheatmap::pheatmap*), as well as score distribution plots (*plotScoreDistribution*).

#### Marker identification and enrichment analysis for the subclusters

Using the *FindMarkers* function in Seurat, markers associated with the subcluster of the enteric neuronal and glial populations were identified. The analysis was limited to genes expressed in at least 25% of cells (min.pct = 0.25) and exhibiting a minimum log fold-change (logFC) of 1.0 between groups. Only genes with an adjusted p-value < 0.05 were considered significant markers. Enrichment analysis was conducted using Gene Ontology (GO), KEGG, and Reactome pathway databases using clusterProfiler (v4.10.1). Enrichment terms with a Benjamini-Hochberg corrected p-value< 0.05 were considered significant. Enrichment results were visualized using ggplot2 or the *sc*.*pl*.*dotplot* function from scanpy (v1.11).

#### Differential expression between conditions

Differential expression analysis between the SNCA 3x and Iso conditions was performed using the *FindMarkers* function in Seurat, with a minimum of 25% cells expressing a given gene (min.pct = 0.25) and a minimum absolute log fold-change threshold of 1. Significant genes were selected based on an adjusted p-value < 0.05, calculated using a Wilcoxon Rank Sum test followed by Bonferroni correction. We used ComplexHeatmap (v2.18.0) to display significant mitochondria-related DEGs across the enteric neuron and glia subclusters.

#### Compositional analysis

Compositional analysis was performed using the scProportionTest package^84^ (v0.0.0.9000), starting with the metadata containing the subcluster and condition information for enteric neurons and glia. The *sc_utils* function was applied to process the metadata for both datasets, and permutation testing was then conducted to evaluate whether the proportions of cells in specific clusters differed significantly between the two conditions (Iso vs. SNCA 3x). The analysis was run with a false discovery rate (FDR) threshold of 0.05 and a log2 fold-change threshold of 1.0.

#### Intercellular communication analysis

Cell-cell communication analysis was performed using the CellChat package^45^ (v2.1.2) to investigate signaling interactions among proliferating cells, enteric neurons and glia, as well as interactions within the subclusters of the latter two populations. Data from both Iso and SNCA 3x conditions were processed individually. For each condition, the dataset was imported into a CellChat object, followed by standard preprocessing steps including normalization and the identification of overexpressed genes and receptors. Communication probability and cellular communication network were then inferred by calculating the probabilities of ligand–receptor interactions based on cell-specific gene expression profiles. Following network aggregation, a merged CellChat object was generated to compare communication differences between Iso and SNCA 3x. Additionally, number of interactions and interaction strength along with the top-ranked signaling pathways were identified and plotted for visualization.

#### Pseudotime analysis of mitochondrial-linked genes

Trajectory analysis of enteric neurons and glia populations was conducted using Slingshot^41^ (v2.10.0). Lineages and pseudotime trajectories were inferred with the *sling-shot* function, specifying the root cluster based on entropy analysis (TSCAN package v1.40.1). Mitochondrial marker genes (*DNM1L, COX4I1, ATP5F1B, NDUFA9, PPARGC1A, PINK1*) were analyzed in enteric neurons and glia by assessing their expression across pseudotime in Iso and SNCA 3x conditions. Pseudotime values were extracted using *sling-Pseudotime()*, averaged across lineages, and validated against cell counts. Linear regression (*lm(expression∼pseudotime * group)*) was performed to assess interactions between pseudotime and condition, followed by FDR correction of p-values to analyze the statistical differences in mitochondrial associated genes expression dynamics between conditions.

#### Metabolic flux analysis

We used COMPASS^85^ (v0.9.10.2) to integrate scRNA-seq data with flux balance analysis, inferring metabolic states in SNCA 3x and Iso enteric neurons and glial cells to compare their metabolic differences. Using a genome-scale metabolic model (GSMM), COMPASS allowed us to integrate transcriptomics data with metabolic network constraints to compute “potential activity” scores for thousands of metabolic reactions across single cells. These scores quantify the degree to which the transcriptomics profiles support flux through specific metabolic reactions. We compared the estimated Log2Fold changes of the metabolic reactions for specific pathways and plotted the difference in Log2Fold values to visualize the changes between reaction scores in SNCA 3x vs. Iso cells. We computed the logarithm of the Wilcoxon adjusted p-value to filter reaction differences. A threshold of 1.3 (corresponding to an adjusted p-value of 0.05) was used to identify significant reactions, where values above this threshold indicated evidence of differential activity between groups.

### ELISA for total and aggregated α-syn

ELISA was used to assess the intracellular levels of total and aggregated α-syn in iPSC-ENLs. Protein was extracted with the buffer provided by the α-syn ELISA kit (cat: ab260052, Abcam) and quantified using BCA assay (Thermo Fisher) for normalization of the data. We used 50mg/mL of extracted protein for the total α-syn analysis and 300ug/mL for the aggregated α-syn (cat: 448807, Biolegend) analyzes. Reactions were performed according to the manufacturer’s instructions, with data analyzed in triplicates.

### Cell viability, Mitotracker and MitoSox assays

Cell viability, mitotracker and mitosox assays were performed using flow cytometry. First, to analyze cell viability we used LIVE/DEAD staining (0.2% in FACS buffer) (Thermo Fisher). To analyze mitochondrial function, we employed 100nM Mitotracker Green (Thermo Fisher) and 50nM Mitotracker Deep Red (Thermo Fisher) to assess percentages of functional and dysfunctional mitochondria in live cells, as described^86^. To assess superoxide production, we employed 5µM MitoSox (Thermo Fisher). ENLs were initially treated with Accutase (Gibco, USA), washed with 1× DPBS, centrifuged at 300g at room temperature for 5 min and then resuspended in FC buffer (2% FCS, 0.01% sodium azide in DPBS). We then counted 500,000 cells per line and proceeded with staining with the probes for 30 min at 37°C and 5% CO2. Cells were then centrifuged, washed twice and resuspended in 300µL FC buffer. At least 30,000 events per sample were analyzed. Data acquisition was performed using a Cytoflex S cytometer (Beckman Coulter) and analysis with the CytExpert 2.4 software.

### Dot-blotting analysis

Cell lysates were diluted in ddH2O (Sigma) to achieve 8µg of total protein in a 3µL drop, which was then spotted onto a nitrocellulose membrane and air-dried for at least 4 hours. Blocking was performed with 5% non-fat dry milk in TBS for 1 hour at room temperature, followed by incubation with the primary antibody (Syn-1, BD Biosciences, cat: 610787, 1:1000) in 5% non-fat milk in 0.1% TBS-Tween20 overnight at 4°C. After three washes with 0.1% TBS-Tween20, membranes were incubated with an HRP-conjugated secondary anti-mouse antibody (Cell signaling, cat: 7076P2, 1:5000) in 5% non-fat milk in 0.1% TBS-Tween20 for 1 hour at room temperature. The blot was then washed three times with 0.1% TBS-Tween20 before signal detection using a chemiluminescence system (ChemiDoc MP, Biorad). Data was normalized to total protein per membrane spot analyzed.

### Fractionation protocol for the enrichment of detergent-insoluble protein aggregates

Detergent-insoluble protein aggregates were enriched using a sarkosyl-based fractionation protocol^56^. Samples were homogenized in ice-cold low salt (LS) buffer (50mM HEPES pH 7.0, 250mM sucrose, 1mM EDTA) with 1X protease and phosphatase inhibitors on ice. Sarkosyl (1% w/v) and NaCl (0.5M) were added, followed by incubation on ice for 15 minutes and sonication with Diagenode Bioruptor Pica (settings: 5s ON, 30s OFF, 5 cycles, medium frequency). Protein concentrations were determined using the BCA assay, and homogenates were ultracentrifuged at 180,000g for 30 minutes at 4°C. Sarkosyl-soluble supernatants were collected, while insoluble pellets were washed with sark-buffer and solubilized in urea buffer (50mM Tris-HCl pH 8.5, 8M urea, 2% SDS) for 30 minutes at room temperature, followed by low-amplitude sonication (setting: 5s ON, 30s OFF, 3 cycles, low frequency). Protein concentrations were normalized based on BCA assay results, and samples were adjusted using sark-buffer before mixing with 5X Laemmli buffer and heating at 95°C for 5 minutes.

#### Western-blotting

For Western blot analysis, proteins were resolved via SDS-PAGE and transferred onto a membrane using wet transfer. Membranes were blocked with 5% BSA in TBS for 1 hour at room temperature, then incubated overnight at 4°C with primary antibodies (mouse anti-α-syn 2A7, Novus, cat: NBP1-05194 and rabbit anti-β-actin, Abcam, cat: ab8227, both 1:1000). Following three washes in 0.1% TBS-Tween20, membranes were incubated with anti-mouse (Cell Signaling, cat: 7076P2) and anti-rabbit (Cell Signaling, cat: 7074S) HRP-conjugated secondary antibodies (both 1:5000) for 1 hour at room temperature. After final washes, protein bands were detected via chemiluminescence (iBright system, Thermo Fisher). Data was normalized to beta-actin.

### Human Cytokine Array

The supernatant of iPSC-ENLs stimulated for 24h with 100ng/mL TNF-α was collected. We added 1mL of the supernatant plus 500µL of array buffer 5 for each membrane of the Proteome Profiler Human Cytokine Array Kit (R&D systems, cat: ARY005B), which detects up to 36 cytokines at the same time. Each membrane was used for the supernatant coming from one biological sample. Analysis was performed according to the manufacturer’s instructions.

### Multielectrode Array recording

To measure neuronal activity, iPSC-ENLs were seeded onto a 48-well CytoView MEA plate (Axion Biosystems) precoated with geltrex on day 15 of differentiation. Cells were maintained in ENC medium until day 70 (end of differentiation). Recordings were performed for 10 minutes at 37°C in an environment of 95% O2 and 5% CO2 using the Maestro MEA System and AxIS software (Axion Biosystems). Signals were sampled at 12.5 kHz with a hardware frequency bandwidth of 200–5000 Hz and later processed using a first-order Butterworth band-pass filter (200–5000 Hz). Spike detection was set at an adaptive threshold of 5.5 standard deviations above background per electrode, with active electrodes defined as those detecting ≥ 5 spikes per minute. Activity parameters were analyzed using the AxIS Metric Plotting Tool (Axion Biosystems).

### Proteomics analysis

Proteomics data were analyzed as previously published, with minor modifications^87^. The cells were lysed with lysis buffer (8M urea, 0.1M NH4HCO3) and the concentration of total proteins was quantified using the BCA kit (Thermo Fisher). Samples were treated with 12 mM DTT for 30 min at room temperature followed by alkylation with 40 mM Iodoacetamide in the dark for 45 min. After dilution of the sample to 1:5 with 0.1M NH4HCO3 an overnight digestion with 1 µg Trypsin (Promega) was performed. The tryptic digest was acidified to pH<3 using TFA and desalted using C18 reversed phase spin columns (Harvard Apparatus) according to the manufacturer’s protocol. Dried peptides were resolved in 0.1% formic acid containing 1% acetonitrile prior to injection into the mass-spectrometer.

The peptide samples were analysed on an Orbitrap Exploris480 mass spectrometer (Thermo Scientific) equipped with a Waters M-class UPLC system (Waters AG) operating in trap/elute mode. Peptides were loaded and chromatographically separated using a Symmetry C18 trap column (5 µm, 180 µm X 20 mm, Waters AG) and as separation column an HSS T3 C18 reverse-phase column (1.8 µm, 75 µm X 250 mm, Waters AG). The columns were equilibrated with 99% solvent A (0.1% formic acid (FA) in water) and 1% solvent B (0.1% FA in ACN). Trapping of peptides was performed at 15 µl/min for 30 sec and afterwards the peptides were eluted using the following gradient: 1-40% B in 120 min; 40-98% B in 5 min. The flow rate was constant 0.3 µl/min and the temperature was controlled at 50°C. High accuracy mass spectra were acquired with a Orbitrap Exploris480 operated in data independent acquisition mode. A survey scan was followed by up to 12 MS2 scans. The survey scan was recorded using quadrupole transmission in the mass range of 350-1500 m/z with an AGC target of 3E6, a resolution of 120’000 at 200 m/z and a maximum injection time of 50 ms. All fragment mass spectra were recorded with a resolution of 30’000 at 200 m/z, an AGC target value of 1E5 and a maximum injection time of 50 ms. The normalized collision energy was set to 28%. Dynamic exclusion was activated and set to 30 sec. For protein identification and label-free quantification the obtained spectra were analyzed using the software FragPipe. This software was used to automatically generate a spectral library and with this the .Raw files were uploaded as input to calculate protein levels.

Differential expression analysis was performed using the limma package, with linear models and contrasts applied to compare Iso and SNCA 3x groups under various treatment conditions. Differentially expressed proteins (DEPs) were identified by filtering for p-values < 0.05 and log2 fold changes ≥ 0.8 or ≤ − 0.8. Data visualization included volcano plots to represent significance and fold change, and Venn diagrams to identify overlapping DEPs across conditions. Gene Ontology (GO) enrichment analysis was performed on DEPs using the clusterProfiler package, focusing on the Molecular Function category, and the results were visualized in bar plots. Final results, including protein identifiers, fold changes, p-values, and GO enrichment, were compiled for downstream analysis. Ingenuity pathway analysis was used to merge the top protein-protein interaction networks found when comparing SNCA 3x vs Iso ENLs stimulated with TNF-α vs basal level.

### Metabolomics analysis

Untargeted metabolomics were in general performed as previously described with minor modifications^88^. Cells were lysed in an 80:20 methanol-water mixture (1 mL per sample) containing recovery standards, followed by centrifugation at 24,000 g and 4°C for 5 min. From the supernatant 350 µL each were used for HILIC and RP chromatography and 230 µL for pooled QC samples, which were prepared the same way as unknown samples after pooling. All supernatants were dried under nitrogen at 30°C before reconstitution in HILIC or RP eluent. LC-MS analysis was performed using a Dionex Ultimate 3000 system coupled to a Q Exactive Focus mass spectrometer (Thermo Fisher Scientific) with HILIC (Acquity UPLC BEH Amide, 1.7 µm) and RP (Acquity UPLC BEH C18, 1.7 µm) chromatography. The system was operated in positive and negative ionization modes with a mass detection range from 66.7 to 1000 m/z in fullscan mode, with the three most abundant features fragmented in ddMS2 mode. QC samples and blanks were included for system calibration and batch correction. Data were analyzed using Compound Discoverer 3.3, with features identified using a 5 ppm mass tolerance and 0.2 min RT tolerance. Peak areas were normalized to protein concentration, Compound annotations were assigned via ChemSpider, mzCloud™, mz-Vault, and in-house databases. Only compounds with annotation levels 1 or 2 (See Table S9) were included. Analysis of individual metabolites was calculated by performing a scaling of the data to consider the donor effect, followed by Student’s unpaired two-tailed t-tests, similar to previous publications^89^. Metabolite-set enrichment analysis was performed using MetaboAnalyst^90^, considering the DEMs (p-values < 0.05, log2 fold changes ≥ 0.5 or ≤ −0.5).

### Integrated pathway analysis

Integrated pathway analysis was generated using MetaboAnalyst^90^ by adding as an input the DEPs (p-values < 0.05 and log2 fold changes ≥ 0.8 or ≤ − 0.8) and DEMs (p-values < 0.05 and log2 fold changes ≥ 0.5 or ≤ − 0.5) obtained when comparing SNCA 3x vs Iso stimulated with TNF-α.

### Proximity ligation assay (PLA)

Imaging 24-well cell plates containing iPSC-ENLs at day 70 (basal and TNF-stimulated) were fixed with 4% paraformaldehyde and permeabilized with 0.4% (v/v) Triton X-100. The Duolink In Situ PLA kit (Sigma Aldrich, cat: DUO92101) was used for PLA. Cells were blocked and incubated with mouse anti-α-syn (Novus, cat: NBP1-05194, 1:50) or rabbit anti-TOM20 (Proteintech, cat: 11802-1-AP, 1:250) antibodies and thereafter with the corresponding PLA probes. After ligation and amplification using the kit reagents, cells were counterstained for α-syn and TOM20 and mounted using Duolink mounting medium with DAPI (1:2500). A pipeline to analyze the PLA dots in the cytoplasm of the cells was created using CellProfiler (Broad Institute) and at least 4-5 individual images per individual biological sample under each condition (basal or TNF) were analyzed. Imaging was performed using the Super Resolution Evident SR Spinning Disk microscope (Olympus) and the cellSens Software and processing was performed using FIJI (ImageJ).

### Mitochondrial parameter analysis

Mitochondrial morphology was assessed using the TOM20 immunofluorescence stainings and analyzed in ImageJ (Fiji) with the Mitochondria Analyzer plugin^91^. Images were acquired as TIFF files, and the TOM20 channel was processed for analysis. Briefly, images were converted to 8-bit grayscale, and background was subtracted using a rolling ball algorithm (radius = 50 pixels). A global threshold (Otsu method) was applied to segment mitochondria, followed by binary processing to refine structures. Nuclei were identified from the DAPI channel using CellProfiler, and mitochondrial parameters were normalized to the number of nuclei per image. A minimum of three images per biological replicate was analyzed, and results were averaged per biological replicate. Statistical analyses were performed in R, with group comparisons conducted using Student’s unpaired two-tailed t-tests.

### Isolation of mitochondrial fractions

Cells were treated with Accutase (Gibco, USA), washed with 1× DPBS, centrifuged at 300g at room temperature for 5min and isolation of mitochondrial fractions was performed using the Mitochondria Isolation Kit for Cultured Cells (Thermo Fisher, cat: 89874), following the manufacturer’s instructions. Isolation was validated via Western blotting using TOM20 (Proteintech, cat: 11802-1-AP, 1:1000) as a mitochondrial marker and β-III-tubulin (Biolegend, cat: Cat:801202, 1:1000) as a cytosolic marker. Proteins were transferred to a membrane using a semi-dry transfer system, followed by blocking and incubation with anti-mouse (Cell Signaling, cat: 7076P2) and anti-rabbit (Cell Signaling, cat: 7074S) HRP-conjugated secondary antibodies (both 1:5000). Bands were visualized using chemiluminescence detection (ChemiDoc MP, Biorad).

### Seahorse analyzes

For Seahorse assays, we seeded 60,000 ENL precursors per cm^2^ on Geltrex (Thermo Fisher)-coated Seahorse XF24 cell plates (Agilent, USA) at day 15 of the differentiation and kept them with ENC medium until day 70, when the analyzes were performed. For mitostress assays, cells were washed two times and incubated for 1h at 37°C in an incubator without CO2, with respiratory medium consisting of DMEM XF (Agilent) supplemented with 10mM glucose solution (Agilent) plus 1mM pyruvate solution (Agilent) and 2mM glutamine (Agilent). After equipment calibration, baseline respiration measurements were followed by injection of 5µM oligomycin (Sigma), 5µM carbonyl cyanide mchlorophenylhydrazone (CCCP) (Sigma) and 5µM rotenone (Sigma) plus 5µM antimycin A (Sigma). To perform the mitofuel assays, we used the same medium and conditions of the mitostress assay, but the compounds were the ones provided by the Seahorse XF Mito Fuel Flex Test Kit (Agilent), in the concentrations indicated for the metabolic dependency analysis, which included 3µM BPTES, 4µM etomoxir and 2µM UK5099 (final concentrations). All respiratory modulators were previously titrated, and all plates were normalized to protein content using the BCA kit (Thermo). Analysis was performed using the Wave software (Agilent).

### Quantification and statistical analysis

Statistical analysis was performed using Graphpad Prism 10, by either two-tailed unpaired Student’s t test (two groups), one-way ANOVA followed by Tukey’s post-hoc test or twoway ANOVA followed by Sidak’s, Tukey’s or Bonferroni’s post hoc test (more than two groups). Error bars represent standard error of the mean (SEM). Statistical analysis for scRNAseq, metabolomics and proteomics analysis were performed by different packages in R as stated. p < 0.05 was considered significant. Details of statistics and sample sizes are described in the figure legends.

## Supplementary Material

### Supplemmentary Figures

The supplementary figures S1-S4 can be found below. Their respective legends are on page 30 of this manuscript.

### Supplementary Tables

The supplementary tables can be found on bioRxiv. Their description is below:

**Table S1**. Primers sequences used in this study

**Table S2**. Subcluster markers annotation, related to figures 2, S2

**Table S3**. SingleR annotation scores, related to figures 2, S2

**Table S4**. Differentially expressed genes between conditions, related to figure 2

**Table S5**. Integrated enrichment of subclusters, related to figure 2

**Table S6**. Cell chat analysis results, related to figures S1, S3

**Table S7**. Pre-processed proteomics data and DEPs, related to figure 5

**Table S8**. Molecular function enrichment of proteomics data, related to figure 5

**Table S9**. Processed metabolomics data, related to figure 5

## Supplementary Figure Legends

**Figure S1.**
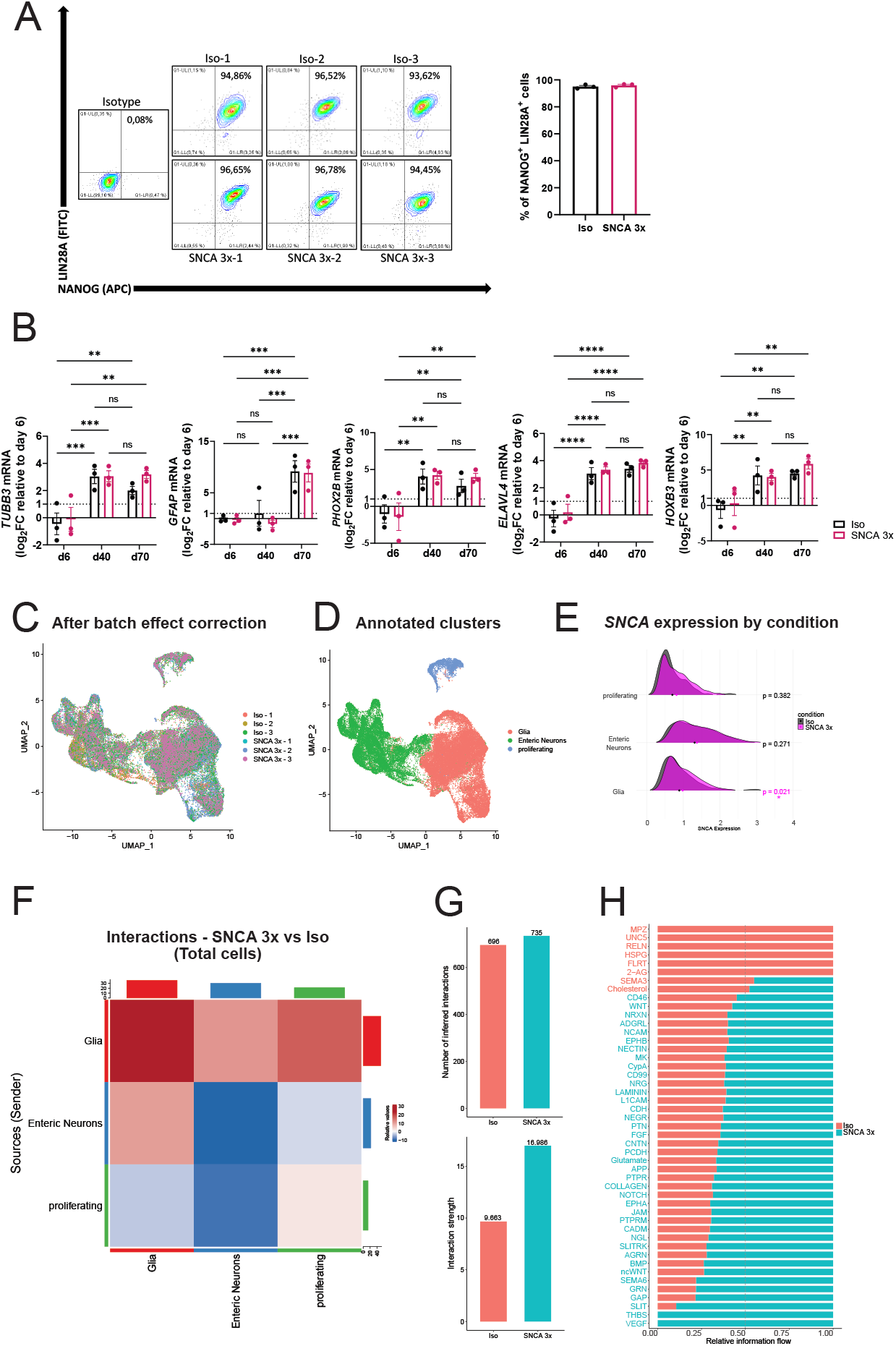
iPSC-ENLs as a model for studying synucleinopathies, related to figure 1. **(A)** Flow cytometry characterization of iPSC purity by staining of NANOG^+^/LIN28A^+^ cells. Analysis was performed before starting the initial timepoint of differentiation (day 0). n=3 biological replicates per group, mean ± SEM. **(B)** Statistical quantification of the gene expression of neuronal marker *TUBB3*, glial marker *GFAP* and enteric neuronal markers *PHOX2B, ELAVL4* and *HOXB3* in iPSC-derived ENLs at days 6, 40 and 70 of differentiation analyzed using RT-qPCR. Log2 fold change was calculated in relation to the Iso group at day 6 of differentiation. n=3 biological replicates per group, mean ± SEM, **p<0.01, ***p<0.001, ****p<0.0001, by two-way ANOVA with Tukey’s post-hoc. **(C)** UMAP plot obtained from scRNA-seq analysis of Iso and SNCA 3x ENLs at day 70 after start of differentiation, showing clustering of each cell line after batch effect correction. **(D)** UMAP plot obtained from scRNA-seq analysis of Iso and SNCA 3x ENLs at day 70 after start of differentiation, showing clusters identified after annotation. **(E)** Ridgeplot showing the expression of SNCA per condition considering each cluster identified. n=3 biological replicates per group, p values calculated by unpaired two-tailed Student’s t test. **(F)** Heatmap comparing the cellular communication between SNCA 3x and Iso ENLs in total cells, with the top color bar representing the sum of the column values displayed in incoming signals and the right color bar representing the sum of outgoing signals, red or blue indicating increased or decreased signal of SNCA 3x compared with Iso, respectively. Data was generated using CellChat. **(G)** Barplots showing the quantification of the number of inferred interactions (top) and interaction strength (bottom) in iPSC-ENLs total cells. Data was generated using CellChat. **(H)** Differences in the overall signaling pathway between SNCA 3x and Iso ENLs in total cells, with the ranking indicating the importance of the pathways; red indicating the signaling pathways enriched in Iso, blue representing the signaling pathways enriched SNCA 3x, and black representing no difference in signaling pathway enrichment in groups.

**Figure S2.**
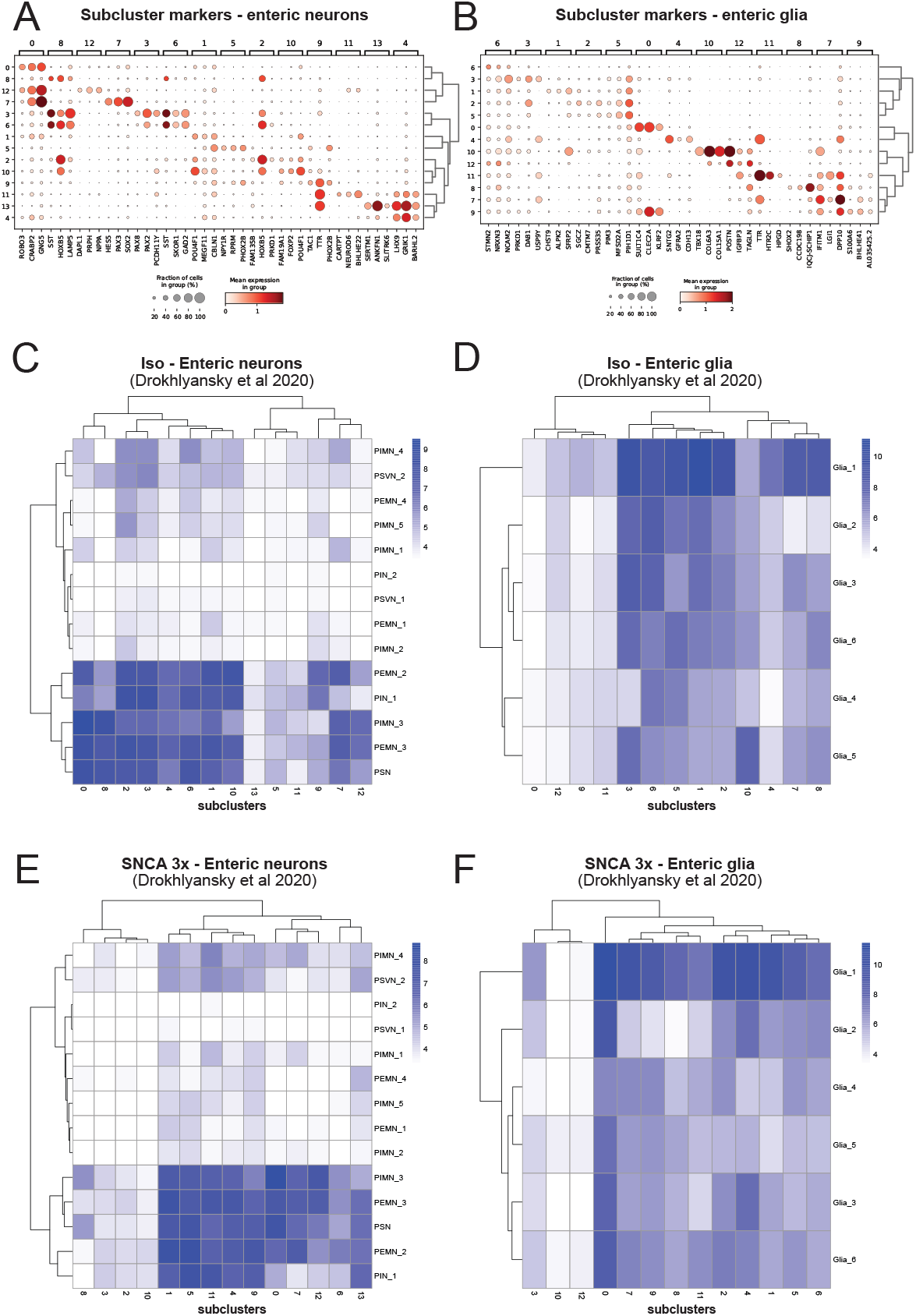
Subclustering identifies distinct iPSC-ENL subpopulations recapitulating the human ENS, related to figure 2. **(A and B)** Dotplots showing the top 3 subcluster markers for enteric neurons (A) and enteric glia (B), with clusterization based on the similarities between subclusters. **(C-F)** Heatmaps showing sublcuster annotation similarities to the human ENS based on singleR scores. C and E represent Iso and SNCA 3x enteric neurons, respectively whereas D and F represent Iso and SNCA 3x enteric glia, respectively. Scores were calculated based on the populations depicted in Drokhlyansky et al 2020^26^.

**Figure S3.**
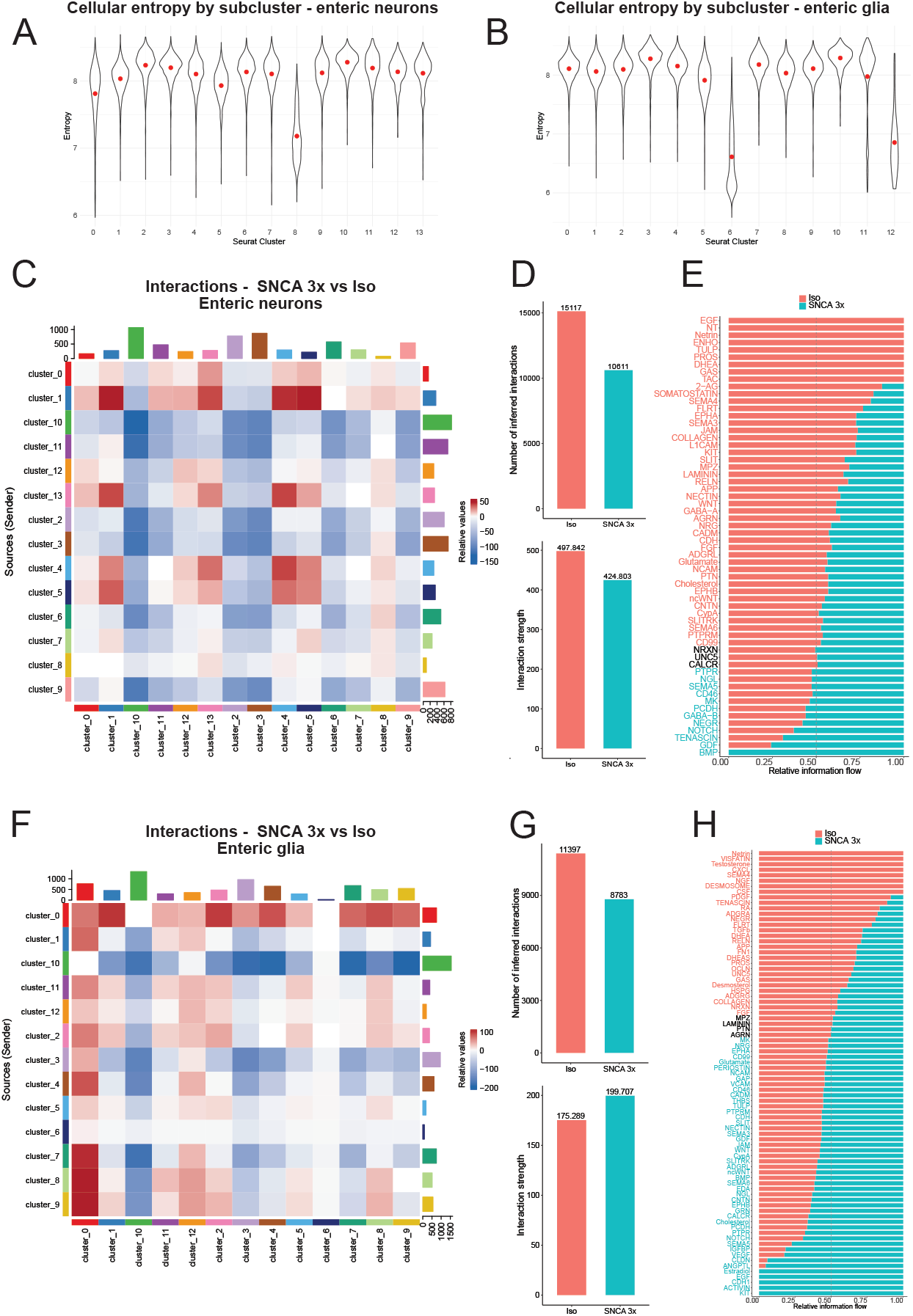
Altered cellular communication and compensatory glial responses to mitochondrial stress in SNCA 3x ENLs, related to figures 2 and 3. **(A and B)** Calculation of cellular entropy by subcluster using TSCAN for enteric neurons (A) and enteric glial cells (B). **(C)** Heatmap comparing the cellular communication between SNCA 3x and Iso ENLs in enteric neuron subclusters, with the top color bar representing the sum of the column values displayed in incoming signals and the right color bar representing the sum of outgoing signals, red or blue indicating increased or decreased signal of SNCA 3x compared with Iso, respectively. Data was generated using CellChat. **(D)** Barplots showing the quantification of the number of inferred interactions (top) and interaction strength (bottom) in enteric neuron subclusters. Data was generated using CellChat. **(E)** Differences in the overall signaling pathway between SNCA 3x and Iso ENLs in enteric neuron subclusters, with the ranking indicating the importance of the pathways; red indicating the signaling pathways enriched in Iso, blue representing the signaling pathways enriched SNCA 3x, and black representing no difference in signaling pathway enrichment in groups. **(F)** Heatmap comparing the cellular communication between SNCA 3x and Iso ENLs in enteric glia subclusters, with the top color bar representing the sum of the column values displayed in incoming signals and the right color bar representing the sum of outgoing signals, red or blue indicating increased or decreased signal of SNCA 3x compared with Iso, respectively. Data was generated using CellChat. **(G)** Barplots showing the quantification of the number of inferred interactions (top) and interaction strength (bottom) in enteric glia subclusters. Data was generated using CellChat. **(H)** Differences in the overall signaling pathway between SNCA 3x and Iso ENLs in enteric glia subclusters, with the ranking indicating the importance of the pathways; red indicating the signaling pathways enriched in Iso, blue representing the signaling pathways enriched SNCA 3x, and black representing no difference in signaling pathway enrichment in groups.

**Figure S4.**
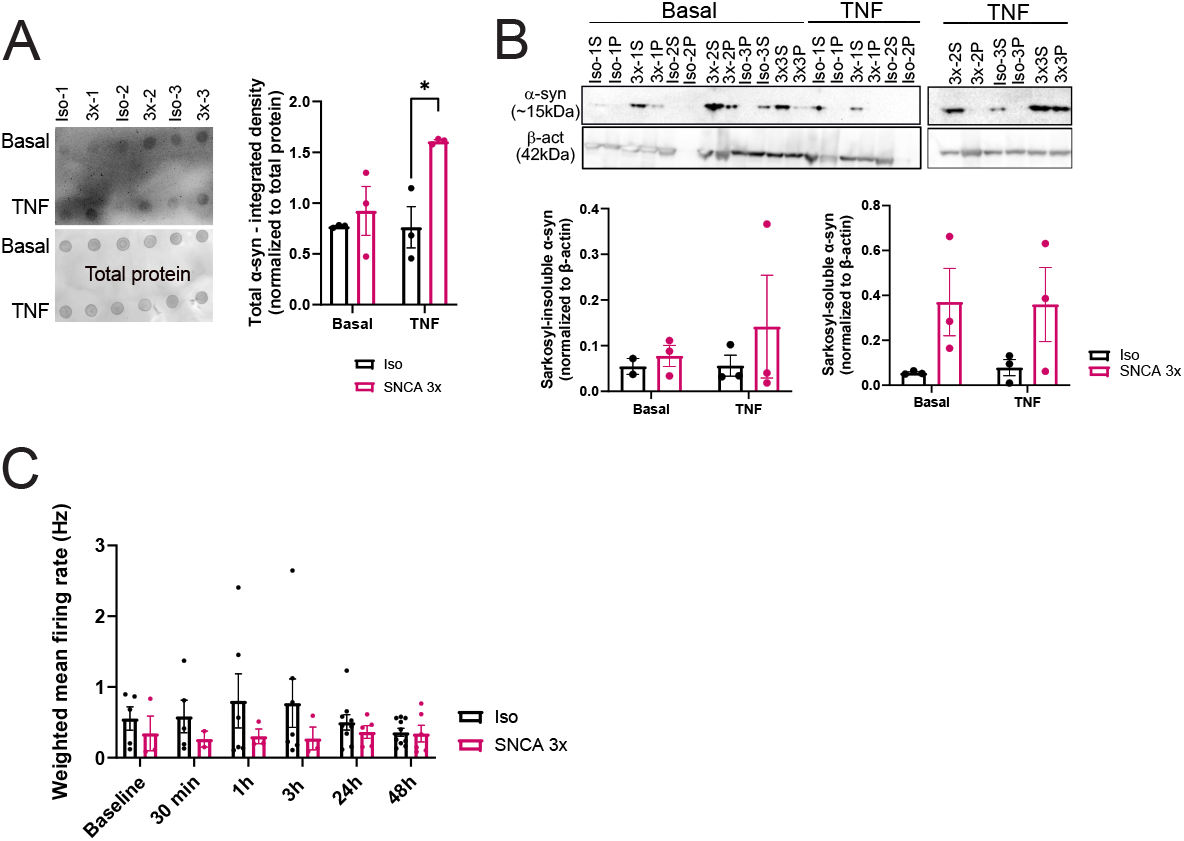
TNF-α uncovers genotype-specific α-syn accumulation and synaptic dysfunction in SNCA 3x ENLs, related to figure 4. **(A)** Dot blot analysis of total α-syn (Syn-1 antibody, BD Biosciences, cat: 610787) quantified intracellularly in iPSC-ENLs, n=3 biological replicates per group, mean ± SEM, *p<0.05 by two-way ANOVA with Bonferroni post-hoc. Basal refers to cells treated with vehicle used to dilute the TNF-α (DPBS+0.1% BSA). **(B)** Sarkosyl-aggregation assay of α-syn blot (2A7 antibody, Novus, Cat:NBP1-05194) and quantification in IPSC-ENLs. Data was normalized to beta-actin. S=supernatant, P=pellet. n=3 biological replicates per group, mean ± SEM. Basal refers to cells treated with vehicle used to dilute the TNF-α (DPBS+0.1% BSA). **(C)** Quantification of the weighted mean firing rate generated from the MEA data, n=wells of a CytoView MEA 48-well plate, representative of 3 biological replicates per group, mean ± SEM.

